# The impact of genetic diversity on gene essentiality within the *E. coli* species

**DOI:** 10.1101/2020.05.25.114553

**Authors:** François Rousset, José Cabezas Caballero, Florence Piastra-Facon, Jesús Fernández-Rodríguez, Olivier Clermont, Erick Denamur, Eduardo P.C. Rocha, David Bikard

## Abstract

Bacteria from the same species can differ widely in their gene content. In *E. coli*, the set of genes shared by all strains, known as the core genome, represents about half the number of genes present in any strain. While recent advances in bacterial genomics have enabled to unravel genes required for fitness in various experimental conditions at the genome scale, most studies have focused on model strains. As a result, the impact of this genetic diversity on core processes of the bacterial cell largely remains to be investigated. Here, we developed a new CRISPR interference platform for high-throughput gene repression that is compatible with most *E. coli* isolates and closely-related species. We applied it to assess the importance of ∼3,400 nearly ubiquitous genes in 3 growth media in 18 representative *E. coli* strains spanning most common phylogroups and lifestyles of the species. Our screens highlighted extensive variations in gene essentiality between strains and conditions. Unlike variations in gene expression level, variations in gene essentiality do not recapitulate the strains’ phylogeny. Investigation of the genetic determinants for these variations highlighted the importance of epistatic interactions with mobile genetic elements. In particular, we showed how mobile genetic elements can trigger the essentiality of core genes that are usually nonessential. This study provides new insights into the evolvability of gene essentiality and argues for the importance of studying various isolates from the same species in bacterial genomics.

## Introduction

Essential genes can be defined as genes required for the reproduction of an organism^1^. The investigation of gene essentiality is key in bacterial genetics for various reasons: (i) in molecular biology, it reveals the most fundamental processes of living cells; (ii) in medical applications, it informs on potential drug targets^2^; (iii) in synthetic biology, it contributes to metabolic engineering efforts^3^ and orients the design of minimal genomes^4^; (iv) in evolutionary biology, it enlightens the gene repertoire of the last universal common ancestor of cellular organisms and provides key phylogenetic markers to establish a tree of Life. Essential genes are thought to be rarely lost because of their essentiality and evolve at a lower rate than other genes^5,6^, a phenomenon likely linked to their higher expression level^7^. However, previous work showed that closely-related taxa have different essential genes^8–14^. Rapid variations in the essential character of a gene might have outstanding implications for the four points raised above.

It is well understood that essential genes depend on growth conditions. Auxotrophic strains lacking genes involved in the synthesis of organic compounds are unable to grow unless the missing compound is available in the medium. The study of single-gene deletion mutant collections such as the Keio collection^15,16^, and transposon-sequencing methods^17–19^ has enabled the determination of genes required for growth in various conditions. However, most studies in *E. coli* were limited to the laboratory-evolved model strain K-12. This strain is not representative of the broad diversity of the *E. coli* species which is characterized by an open pangenome with high rates of horizontal gene transfer (HGT)^20–22^. The pangenome of *E. coli* comprises a large majority of accessory genes (i.e. not present in all strains) while genes present in more than 99% of isolates represent less than 50% of the average gene content^22^. This broad genetic diversity results in the adaptation of *E. coli* strains to multiple ecological niches and lifestyles. *E. coli* can be found in the environment as well as in association with humans and animals where it can behave as a gut commensal or as an opportunistic intestinal and extra-intestinal pathogen^23^. This highlights the importance of studying gene essentiality beyond the strain K-12^24^. A few studies have used transposon-based methods to determine the genetic requirements of clinical *E. coli* isolates for *in vitro* growth or colonization of animal models^25–29^. This showed that clinical strains associated with different pathologies require different genes for colonization and virulence. Although these findings represent an important insight into the mechanisms of infection, a direct comparison of growth requirements of *E. coli* strains is still lacking. In particular, the broad genetic diversity of *E. coli* provides the opportunity to assess how the genetic background influences gene essentiality.

Several reasons could explain why genetic diversity may impact gene essentiality. A gene that is essential in a strain might be dispensable in another strain if the latter carries a homolog or an analog that performs the same function. In this situation, the pair of genes is known as synthetic lethal. Another example is the situation of prophage repressors and antitoxins^30^. These horizontally-transferred elements typically belong to the accessory genome and are only essential when the cognate prophage or toxin is also present. However, it is unclear if there are significant variations in the essential character of core genes across the *E. coli* species. A recent investigation of a panel of 9 *Pseudomonas aeruginosa* strains showed that gene essentiality indeed varies between strains^11^, but the underlying mechanisms and the relevance of these findings to other bacterial species remain to be investigated.

To tackle this question, we turned to CRISPR interference (CRISPRi). This method is based on the catalytically-inactivated variant of Cas9, dCas9, which can be directed by a single-guide RNA (sgRNA) to bind a target gene and silence its expression^31,32^. Genome-wide CRISPRi screens were recently employed in *E. coli*^33–35^ and in a few other bacterial species^36,37^. A custom sgRNA library can be designed to target genes of interest and introduced into a pool of cells. The impact of silencing individual genes on cell growth can then be measured by monitoring the fold-change in the abundance of each sgRNA. Here, we first developed an easy-to-use CRISPRi screening platform compatible with most *E. coli* isolates and closely-related species. We then designed a compact sgRNA library targeting the *E. coli* core genome (here taken as genes present in more than 90% of isolates) in order to compare the essentiality of core genes in different genetic backgrounds and growth conditions. Our results reveal how the essentiality of core genes can substantially vary at the strain level. Further investigation of the underlying mechanisms showed that HGT and gene loss events can modulate the essentiality of core genes.

## Results

### A broad CRISPRi plateform for *Escherichia* and closely-related species

We first designed an easy-to-use single plasmid vector for CRISPRi called pFR56, comprising a constitutively expressed sgRNA, a *dCas9* expression cassette controlled by a DAPG-inducible PhlF promoter and an RP4 origin to enable transfer by conjugation. In order to ensure plasmid stability in most strains, protein-coding sequences from pFR56 were recoded to avoid the restriction sites that are recognized by the most common restriction-modification systems of *E. coli*^38^. In a previous study, we identified a sequence-specific toxicity of dCas9 in *E. coli*, termed the “bad-seed” effect, which can be alleviated by decreasing dCas9 concentration^33^. In order to avoid this toxicity effect, we optimized the dCas9 expression level from pFR56 to maintain a strong on-target repression while ensuring that sgRNAs bearing a bad-seed sequence induce no visible fitness defect (**Supplementary Fig. 1a**). pFR56 achieved a high conjugation rate and was stable for >24 generations without antibiotic selection in various strains from species belonging to the *Escherichia, Klebsiella* and *Citrobacter* genera (**Supplementary Fig. 1b-c**). We tested dCas9-mediated repression with a sgRNA targeting the essential gene *rpsL* in these strains. In all of them, induction of dCas9 expression strongly inhibited growth. (**Supplementary Fig. 1d**). This demonstrates the usefulness of our CRISPRi system in a broad range of *E. coli* isolates and in closely related species.

### A compact sgRNA library targeting ∼3,400 nearly-ubiquitous genes from *E. coli*

While most studies of *E. coli* rely on lab-evolved derivatives of strain K-12, we aimed at investigating gene essentiality in the *E. coli* species as a whole. The size of the pangenome makes it impossible to target all genes from the species. Instead, focusing on persistent genes enables a direct comparison of the same genes under different genetic backgrounds. We analyzed 370 complete *E. coli* genome sequences and identified 3380 protein-coding genes present in > 90 % of genomes (**Fig. 1a**). We then selected 3-4 sgRNAs per gene following rules designed to favor targets that are conserved across strains while minimizing off-target activity and avoiding toxic seed sequences^33^ (**see Methods**). We also used a model of dCas9 on-target activity to select the most active guides^39^. Our library also includes guides targeting rRNAs, tRNAs and widespread ncRNAs. The resulting *E. coli* core genome (EcoCG) library comprises 11,629 sgRNAs targeting ∼60 to 80% of the protein-coding genes of any *E. coli* strain as well as 100% of rRNAs, 75-85% of tRNAs and 15-25% of annotated ncRNAs (**Fig. 1b**). The EcoCG library was cloned into pFR56 and transferred to K-12 MG1655 by conjugation in order to evaluate its performance in the prediction of essential genes. The library was grown in LB supplemented with DAPG to induce dCas9 and re-diluted twice to achieve 20 generations of growth. The pFR56 plasmid was extracted at the beginning and at the end of the experiment followed by sequencing of the library to monitor changes in the frequency of each sgRNA (**Fig. 1c**). Our screen achieved a very high biological reproducibility (Pearson’s r> 0.99) (**Fig. 1d**). For each sgRNA, we calculated the log2-transformed fold change (log2FC) in sgRNA abundance as a measure of sgRNA effect on cell fitness and the median log2FC value was used to score genes. As expected, most depleted sgRNAs corresponded to known essential genes of K-12 MG1655^17^ and our library predicted essential genes better than our previous randomly-designed genome-wide library^34^ (AUC = 0.979 vs 0.963) despite being much smaller (∼9 vs 3.4 sgRNAs per gene on average) (**Fig. 1e**). This is partly explained by the improved library design which results in a lower within-gene standard-deviation of the log2FC values (**Fig. 1f**). Overall, this shows the efficiency of our library to predict essential genes with high confidence.

**Figure 1.**
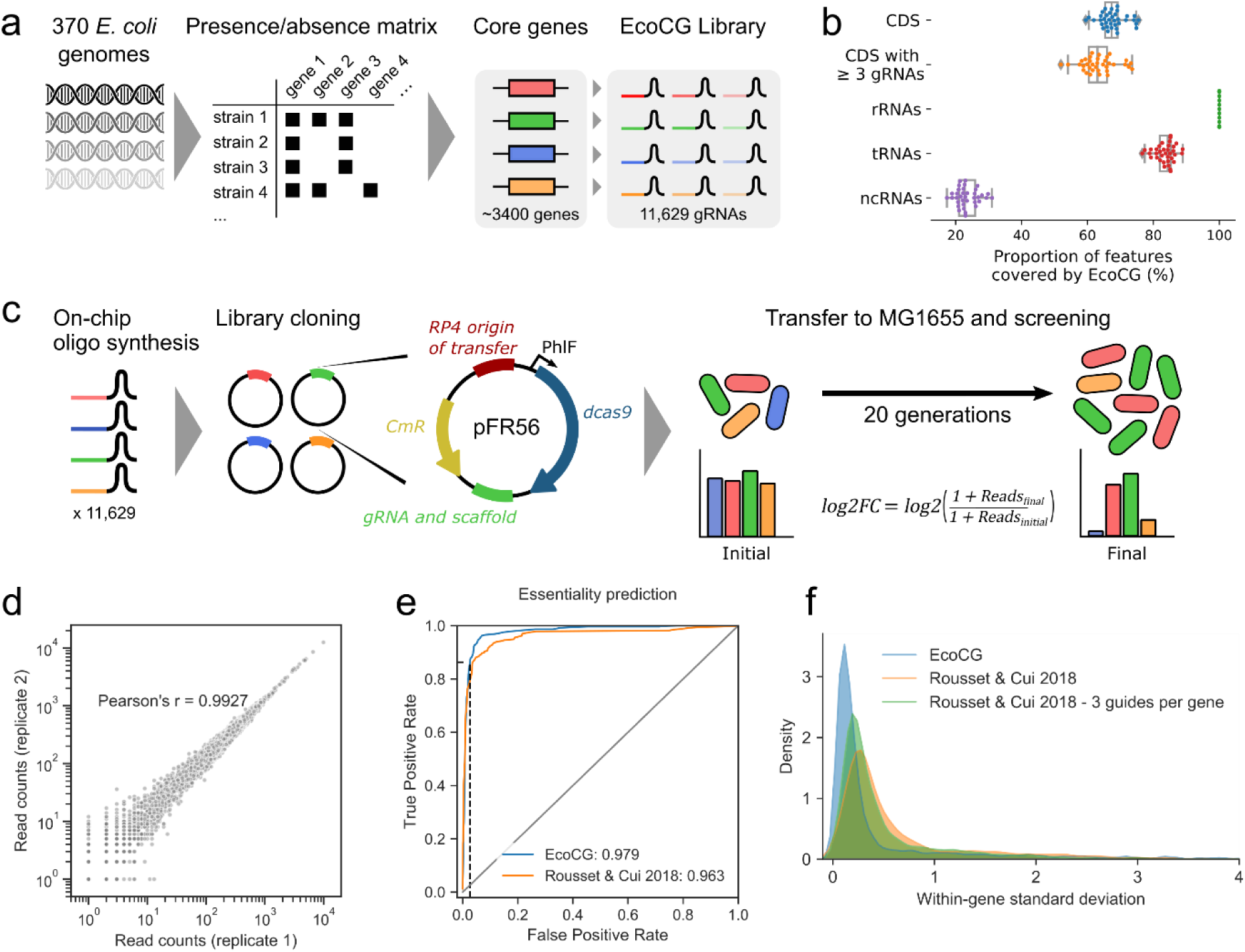
Robust screening of ∼3400 conserved *E. coli* genes with the EcoCG library. **a**, Starting from 370 complete *E. coli* genomes, a gene presence/absence matrix was computed to deduce 3,380 protein-coding genes that are present in > 90% *E. coli* strains. For each gene, 3 or 4 sgRNAs were selected based on the proportion of targeted strains and on the predicted off-target activity, efficiency and bad-seed effect. We also added sgRNAs targeting rRNAs, tRNAs and widespread ncRNAs (see Methods), yielding the EcoCG library comprising 11,629 sgRNAs. **b**, The EcoCG library was mapped to the genome of 42 strains. On average, it targets 67.7% of the protein-coding gene content (with 100% nucleotide identity), and 63.4 % with at least 3 sgRNAs. **c**, The EcoCG library was synthesized and cloned onto pFR56, our single-vector CRISPRi system. The library was transferred to K-12 MG1655 by conjugation and the abundance of guides in the library was monitored by deep sequencing before and after 20 generations of growth with dCas9 induction. **d**, The correlation of experimental replicates suggests an excellent reproducibility. **e**, The log2FC value was calculated for each sgRNA and the median log2FC value was used as a gene score to predict gene essentiality using the TraDIS dataset^17^ as ground truth. The plot shows the receiver operating characteristic (ROC) curve of the prediction model. The dashed black line marks the threshold chosen in further analysis (gene score < −3). **f**, For each gene, the standard-deviation of log2FC values of the different sgRNAs was calculated for the EcoCG library and for our previous genome-wide library^33,34^. In order to account for the difference in library size, we also calculated for each gene the mean standard-deviation of log2FC values obtained from all permutations of 3 sgRNAs from our previous library.

### Distribution of gene essentiality in an *E. coli* strain panel

We selected a panel of 18 *E. coli* isolates from a collection of 92 *E. coli* natural isolates spanning most common *E. coli* phylogroups (A, B1, B2, D, E and F) and lifestyles in order to compare the essentiality of their conserved genes (**Supplementary Table 1, see Methods**). This panel includes the lab-derived strain K-12 MG1655, environmental isolates (E1114, E1167 and E101), commensals from humans (HS) and other mammals (M114, ROAR8, TA054, TA249, TA280 and TA447), an intestinal pathogen associated with Crohn’s disease (41-1Ti9) and extra-intestinal pathogens isolated from blood, lungs, urine and cerebrospinal fluid from humans and poultry (H120, JJ1886, APEC O1, S88, CFT073, UTI89). In order to compare genetic requirements for growth in various experimental contexts, we performed CRISPRi screens with each strain in two biological replicates during aerobic growth in LB or in minimal M9-glucose medium (M9), as well as during anaerobic growth in gut microbiota medium (GMM) (**Supplementary Table 2, Fig. 2a**), yielding a total of 100 CRISPRi screens on ∼3400 genes (four strains were discarded in M9 due to insufficient growth). After 20 generations with dCas9 induction, cells were collected and the library was sequenced. Biological replicates achieved a very high reproducibility (median Pearson’s r = 0.988), demonstrating the robustness of the method (**Supplementary Fig. 2a**). For each screen, we calculated the log2-transformed fold-change (log2FC) and the gene score as described above (**Supplementary Tables 3-4**), resulting in a gene-strain scoring matrix for each of the three tested media. In the following analyses, we considered genes with a score lower than −3 as essential in a given strain and condition. This stringent threshold recovers 86.3% of known essential genes from K-12 in LB^17^ with a false-positive rate of only 2.7% (**Fig. 1e**), mostly due to expected polar effects^34^.

**Figure 2.**
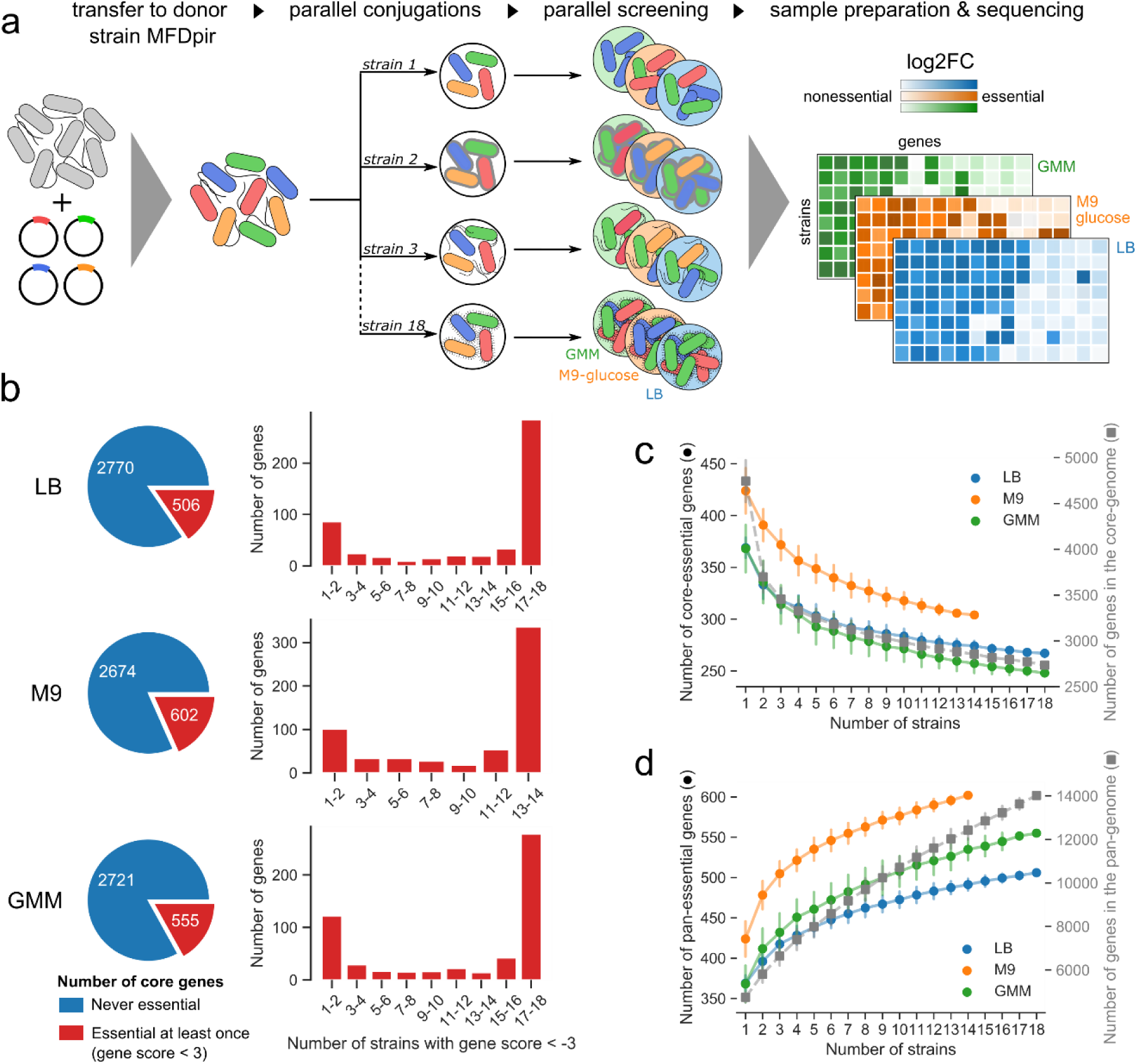
Distribution of fitness defects after CRISPRi screens in 18 *E. coli* strains and 3 media. **a**, The MFDpir conjugation strain^42^ was used to transfer the EcoCG library to a panel of 18 *E. coli* isolates. Each strain was then grown for 20 generations with dCas9 induction in aerobic condition in LB and M9-glucose medium and in anaerobic condition in gut microbiota medium (GMM). Log2FC and gene score values were computed (**see Methods**). Only 14 strains were screened in M9-glucose due to poor growth of 4 strains. **b**, For each medium, we selected core genes whose repression induces a fitness defect in at least one strain (gene score < −3) (left) and reported the number of strains where this defect can be seen (right). **(c-d)** Evolution of the number of core genes that are essential in all strains (**c**) or in at least one strain (**d**) as a function of the number of selected strains (circle markers). The error bars indicate the standard-deviation of up to 250 random permutations. The grey dashed curves represent the size of the core genome (**c**) or the size of the pangenome (**d**) (Square markers) with the scale shown on the right.

We first investigated the overall genetic requirements of the *E. coli* species for growth in each medium by considering the median gene score across all strains. We identified 366 essential protein-coding genes in LB, 427 in M9 and 372 in GMM (median gene score < −3). Genes involved in the biosynthesis of amino acids, nucleotides and cofactors are required in M9 due to the absence of these compounds in the medium. This largely explains the higher number of genes required for growth in M9 compared to LB or GMM (**Supplementary Fig. 3a**). Some genes are specifically nonessential in GMM, in particular genes involved in aerobic respiration (*cydAB, cydX, cydCD*) and TCA cycle (*lpd, icd, acnB*) which are probably attributable to the anaerobic condition of the screen in GMM (**Supplementary Fig. 3b**). Conversely, some genes are specifically essential in GMM such as the fermentation gene *adhE* (alcohol dehydrogenase / aldehyde-dehydrogenase) and the ATP synthase genes. Indeed, in aerobic conditions, the ATP synthase uses the H^+^-gradient to synthetize ATP from ADP, which is not a strictly essential process since ATP can be obtained from other sources. However, in anaerobic conditions, the ATP synthase hydrolyzes ATP to create the H^+^-gradient required for membrane function^40^. Interestingly, other genes that are specifically essential in GMM include the Ribo ome assembly factor *bipA*, the alanine racemase I *alr* and components of the Tol/Pal system. Overall, these results better define the most common genetic requirements of *E. coli* across growth media.

We then explored how many genes are essential in at least one strain and how frequently these genes are essential (**Fig. 2b**). We found many more genes that are essential in at least one strain than genes that are essential in all tested strains (506 vs 267 in LB, 602 vs 364 in M9 and 555 vs 248 in GMM). Most essential genes are either essential in most strains or in a small number of them. This shows that the essentiality of core genes varies substantially at the strain level. We can tentatively use this data to define a core-essential genome, i.e. genes that are virtually essential in all strains of the species, and a pan-essential genome, i.e. genes that are essential in at least one strain of the species. We performed a rarefaction analysis by computing the core-essential genome and pan-essential genome for various sets of strains. Interestingly, the size of the core genome and the size of the core-essential genome converge at a similar pace (**Fig. 2c**). As a result, the fraction of the core genome that is essential in all strains is roughly independent from the number of strains under consideration (e.g. ∼9-10% of the core genome in LB) (**Supplementary Fig. 4a**). The set of core genes that are essential in at least one strain keeps increasing with the addition of new strains (**Fig. 2d, Supplementary Fig. 4b**), showing that our results probably only reveal a fraction of the existing differences at the species level: when all strains are considered, up to 18.5% of their core genome is essential in at least one strain in LB (21.4% in M9 and 20.3% in GMM). This suggests that a significant part of the nonessential core genome is likely to become essential in certain genetic backgrounds.

### Phylogeny explains differences in gene expression profiles but not in gene essentiality

We then tried to gain insight into the factors shaping these differences. Since gene essentiality has been linked to a higher gene expression level^41^, we wondered to what extent changes in gene essentiality are reflected by changes in gene expression level. We generated RNA-sequencing (RNA-seq) data for 16 strains during growth in exponential phase in LB and compared the expression of core genes (**Supplementary Table 5, see Methods**). As previously observed, expression level and essentiality were correlated, with a higher expression level for essential genes (**Supplementary Fig. 5a**). We wondered whether this was also the case for genes whose essentiality varies, i.e. if a shift in essentiality is associated with a shift in expression. We selected 87 genes that were variably essential between the 16 strains assayed in RNA-seq experiments (**see Methods**). Considering all strains together, these “variably essential” genes tend to be more expressed than genes that are never essential but less expressed than genes that are always essential (**Supplementary Fig. 5b**). When considering each “variably essential” gene individually, we found no correlation between CRISPRi fitness and its gene expression level across the 16 strains (**Supplementary Fig. 5c**). This suggests that a shift is essentiality is not associated with a shift in expression level.

We then investigated the relevance of vertical evolution to the variations in gene expression and essentiality. Strikingly, we observed a strong negative correlation between the phylogenetic distance of pairs of strains and the similarity in their gene expression profile (Spearman’s rho = −0.52, p = 10^−9^), i.e. closely-related strains have more similar expression profiles (**Fig. 3a**). Interestingly, K-12 MG1655 seems to be an outlier and discarding it from this analysis markedly improved the correlation (rho = −0.69, p < 10^−15^) (**Fig. 3a and Supplementary Fig. 6a**). This is possibly linked to the high number of mutations acquired during laboratory evolution. We then conducted the same analysis with the CRISPRi fitness profiles. In contrast with the gene expression profiles, the correlation between phylogenetic distance and similarity in gene essentiality was very weak and barely significant in LB (rho = −0.2, p = 0.01) (**Fig. 3b**) and was absent in M9 and GMM (**Supplementary Fig. 6b-c**), regardless of the inclusion of K-12 MG1655. Altogether, this suggests that while changes in the expression level of core genes are strongly linked to vertical inheritance, changes in gene essentiality are not.

**Figure 3.**
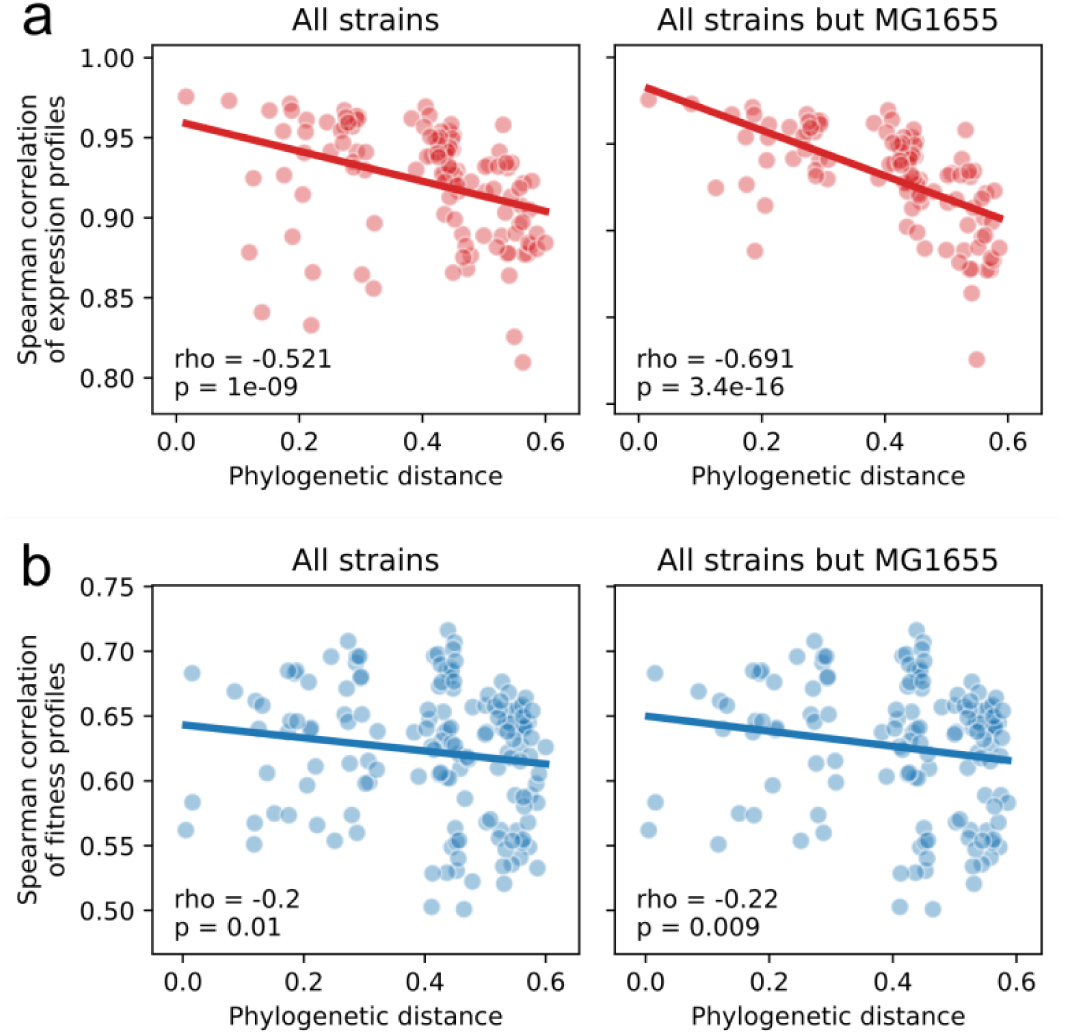
Changes in gene expression but not in gene essentiality recapitulate phylogeny. **a**, RNA-seq data obtained in 16 strains during exponential growth in LB was used to calculate Spearman correlation coefficients of all pairs of gene expression profiles. This showed a strong correlation between the similarity in gene expression and the evolutionary distance of pairs of strains (rho = −0.521, p = 1.10^−9^, N = 120) (left). Removing the lab strain MG1655 from this analysis substantially improved this correlation (rho = −0.691, p < 4.10^−16^, N = 105) (right). **b**, Similarly, pairwise Spearman correlation coefficients in gene essentiality were calculated from CRISPRi screens in LB, showing a weak correlation with phylogenetic distance (rho = −0.2, p = 0.01, N = 153 with MG1655 (left), rho = −0.22, p = 0.009, N = 136 without (right)).

### Homologs rarely provide functional redundancy

In order to investigate the genetic mechanisms that explain the differences in essentiality, we focused on cases where the difference is very strong by selecting genes whose repression induces a strong fitness defect (stringent gene score < −5) in at least one strain, while having no effect (gene score > −1) in at least one strain (**see Methods**). This resulted in 32 protein-coding genes in LB, 55 in M9 and 66 in GMM which displayed very distinct degrees of essentiality across strains (**Fig. 4**). We then tried to determine the genetic basis explaining some of these differences.

**Figure 4.**
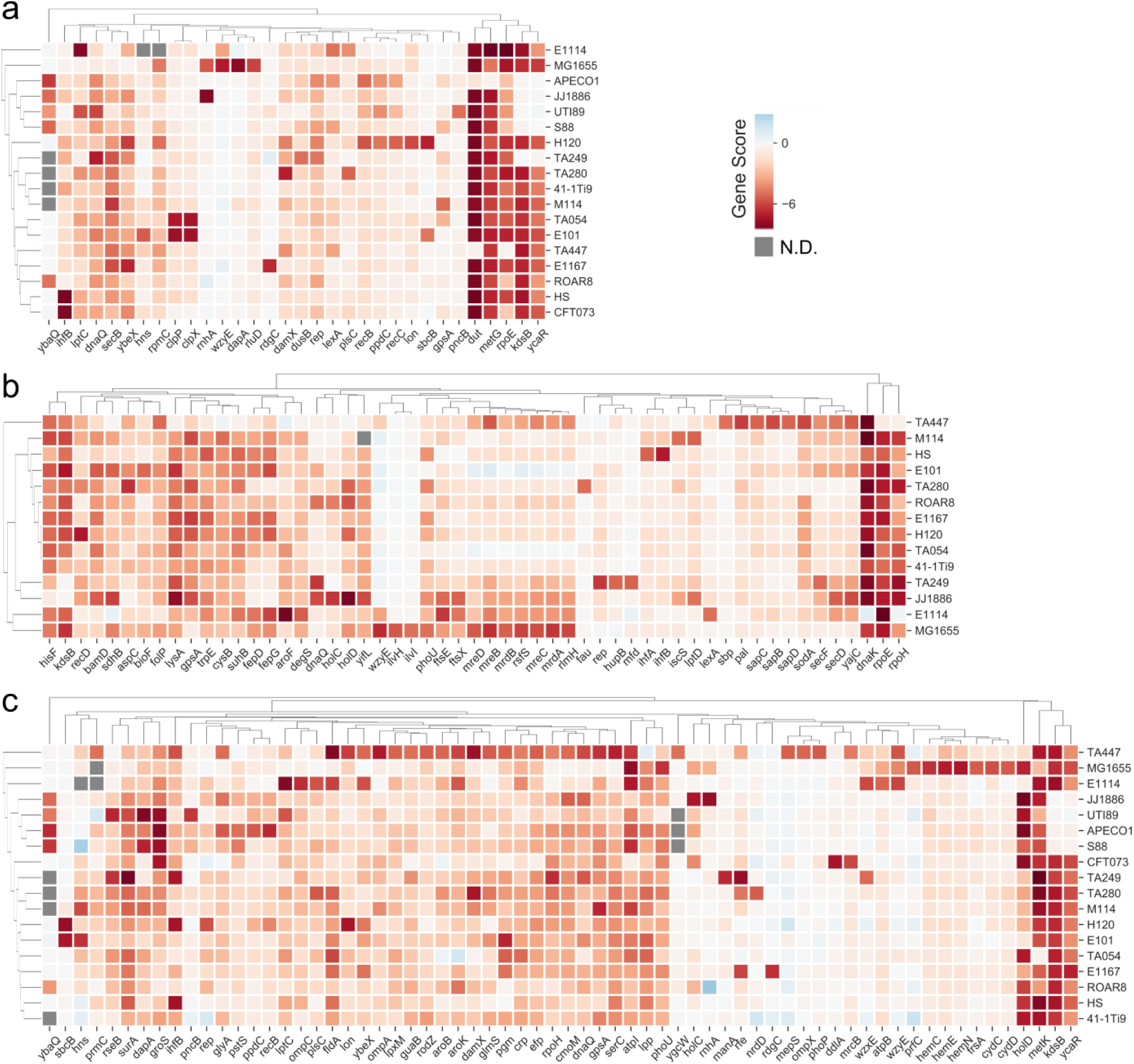
Extensive differences in gene essentiality within *E. coli* core genes. For each experimental condition, we selected genes whose repression produces a strong fitness defect in at least one strain (gene score < −5) while producing no effect in at least one strain (gene score > −1) (see Methods). Panels (**a**), (**b**) and (**c**) show a heatmap of the selected genes in LB, M9 and GMM respectively. Rows and columns were clustered by Euclidian distance. Grey squares correspond to genes and strains where no gene score is available (N.D., non-determined).

A gene that is essential in a genetic background may be lost in another background without loss of cell viability for several reasons. The loss may be rescued (i) by the presence of a homologous or analogous gene of similar function, (ii) by another system performing the same function, or (iii) by epistatic interactions such as in toxin-antitoxin systems or phage repressor genes. We first investigated whether some of the differences in essentiality could be linked to the presence of functional homologs of essential genes in some strains. Our screens showed that all strains where the *ycaR*-*kdsB* transcriptional unit (expressing the essential CMP-KDO synthetase KdsB) is dispensable carry another CMP-KDO synthetase gene, *kpsU*, whose product shares 46% of identity with KdsB. Simultaneous repression of both *kdsB* and *kpsU* induced a strong fitness defect in strains that are resistant to *kdsB* knockdown (**Supplementary Fig. 7**). Interestingly, CFT073 and 41-1Ti9 also carry *kpsU* but remain sensitive to *kdsB* knockdown suggesting that *kpsU* might not be expressed or functional in these strains.

We investigated if a similar mechanism could explain the case of *metG* (methionine-tRNA ligase) which is essential in all strains except APEC O1 (**Fig. 4**). While resequencing the genome of APEC O1, we observed the presence of an unreported plasmid carrying an almost identical copy of *metG*. Hence, the gene is not essential in this strain because its loss is compensated by a close homolog. Interestingly, the same plasmid was also found in one other strain (TA447) where the gene remains essential. This can likely be explained by the fact that sgRNAs present in the library target both copies of *metG* in this strain, while in APEC O1 there are single-nucleotide variants between the two homologs preventing dCas9 from targeting the plasmidic copy of the gene. We could identify another case of genetic redundancy likely missed by our screen because dCas9 targets both copies of the gene: strain TA249 carries two copies of *glnS* (glutamine—tRNA ligase) sharing 96.7% of nucleotide identity. Gene duplications are frequent and transient in bacteria such that duplicates are typically identical^43^. In this context, the ability of dCas9 to silence several copies of a gene means that this duplication phenomenon does not affect the identification of essential genes by our method.

We attempted to assess how frequently the existence of homologs makes an essential gene dispensable. Overall, the strains we tested carry 10 to 17 (median = 13.5) homologs (>40% identity) of core genes that are essential in *E. coli* K-12 in LB^17^, including 3 to 7 (median = 4) with >60% identity. This shows that homologs of essential genes are relatively frequent. We might therefore expect more cases of essential genes that become nonessential in some strains because of genetic redundancy. However, this seems to be the case only for the genes detailed above. As an example, *nrdA* and *nrdB* remain essential in APEC O1 and TA447 despite the presence of two homologs whose product shares 63% and 60% identity to NrdA and NrdB respectively. RNA-seq data showed that these homologs are poorly expressed in our experimental conditions (< 2% of the expression level of *nrdA* and *nrdB*) which likely explains their inability to make *nrdAB* nonessential. In addition, putative homologs of essential genes may not be functionally redundant since sequence homology does not necessarily imply functional redundancy^44^. Altogether, our data shows that with a few exceptions documented here, the presence of a homolog does not provide functional redundancy.

### Functional redundancy provided by a non-homologous gene

A gene may also become non-essential because of the acquisition of genes of analogous function by HGT. The product of *dut* (dUTPase) hydrolyses dUTP into dUMP in order to avoid its incorporation into DNA^45^. This gene is essential except in APEC O1 and TA447 (**Fig. 4**), but we did not find any sequence homolog of Dut in these two genomes. We successfully built a TA447*Δdut* strain and verified the absence of compensatory mutations, confirming that *dut* is indeed nonessential in this strain. Further investigation showed that the plasmid that is shared between APEC O1 and TA447 contains nucleotide biosynthesis genes (**Fig. 5a**). In particular, a hypothetical protein (TA447_03166, accession: WP_085453089.1) that is present in ∼2.6% of available *Escherichia* genomes has an ATP-pyrophosphohydrolase-like domain and shows structural homology to a MazG-like protein from *Deinococcus radiodurans* and other NTP-pyrophosphatases. This gene became essential in TA447 when *dut* was deleted (**Fig. 5b**), and rescued the growth of K-12 MG1655 when *dut* was repressed (**Fig. 5c**). Interestingly, *dut* repression induced a fitness defect in TA447 in GMM (**Fig. 5d**). We showed that in this medium, TA447_03166 is expressed at only ∼60% of its expression level in LB (**Fig. 5e**), which may be insufficient to complement the loss of *dut*. Taken together, these results show that TA447_03166 is probably able to convert dUTP into dUMP. In this case, compensation is provided by a functional analog sharing no sequence homology with the essential protein and the compensation depends on the level of expression of the analog.

**Figure 5.**
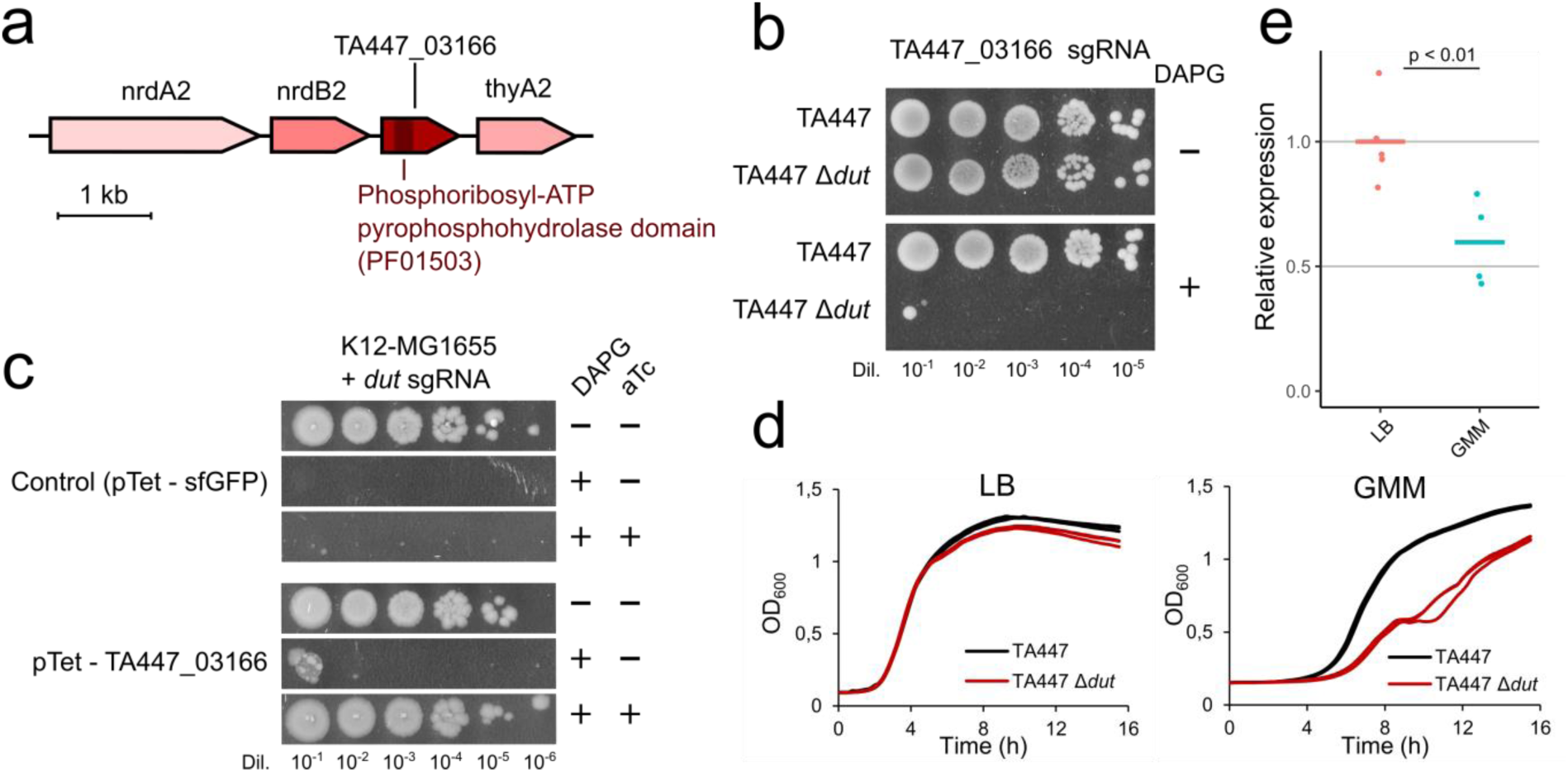
A plasmid-borne dUTPase makes *dut* nonessential in strains TA447 and APECO1. **a**, A region encoding nucleotide biosynthesis genes was identified in TA447 and contains a gene encoding a hypothetical protein (TA447_03166) with a Phosphoribosyl-ATP pyrophosphohydrolase domain (pfam 01503). **b**, Drop assay showing that targeting this gene with dCas9 is lethal when *dut* is deleted from TA447. **c**, This protein was cloned and expressed from an aTc-inducible pTet promoter in K12-MG1655. dCas9-mediated silencing of *dut* has no effect when this protein is expressed. **d**, Growth curves were performed in triplicates in LB or GMM with TA447 and TA447*Δdut*. **e**, The relative expression of TA447_03166 was measured in LB and in GMM. The horizontal bars show the mean of 3 biological replicates and 2 technical replicates.

### A conserved transcriptional regulator controls the expression of a clade-specific toxin

Elements that are essential because they suppress an accessory toxin or phage are typically found in the accessory genome rather than in the core genome. We could nonetheless identify a conserved antitoxin displaying variable essentiality. It is one of the rare cases of a gene essential in an intermediate number of strains (5/18). *ybaQ* encodes a DNA-binding transcriptional regulator of unknown function. The analysis of the genetic organization and conservation of its locus suggests that *ybaQ* is essential in strains from phylogroups B2 and F that carry the *higB-1* toxin upstream *ybaQ*, except in CFT073 where *higB-1* is truncated by a stop codon (**Supplementary Fig. 8a-b**). In these strains, the promoter of *higB-1* contains the YbaQ binding motif^46^ and we found that *ybaQ* knockdown strongly overexpresses *higB-1* in the pathogenic strain S88 (**Supplementary Fig. 8c**). Interestingly, *higB-1* is absent in most phylogroups, including in K-12, while both *higB-1* and *ybaQ* were lost in the phylogroup D. Taken together, these findings suggest that the main role of *ybaQ* was originally to control the expression of the *higB-1* toxin which was then lost in some clades. The conservation of the *ybaQ* antitoxin even in the absence of the cognate toxin suggests that this gene might have other functions that remain to be investigated.

### Epistatic mutations can trigger the essentiality of core genes

In contrast with *kdsB, metG, dut* and *ybaQ*, most “variably essential” genes are essential in very few strains (**Fig. 2b** and **Fig. 4**). This suggests that genes can frequently become essential in a few genetic backgrounds. Most strains (72%, 13/18) had at least one strain-specific or near-specific (in ≤ 2 strains) essential gene in at least one medium and K-12 MG1655-specific essential genes were the most abundant. We first looked for evidence explaining such differences in the literature. For instance, *rluD* (23S rRNA pseudouridine^1911/1915/1917^ synthase) is known to be essential in K-12 because of an epistatic interaction with a mutation acquired in *prfB* during laboratory evolution that is absent in other strains^47^. Other cases of epistatic interactions include the acetohydroxy acid synthase III (AHAS III) encoded by *ilvHI* in K-12 in M9 medium (**Fig. 4b**). An isozyme encoded by *ilvGM* can perform the same reaction in other strains but is disrupted in K-12 by a frame-shift, making AHAS III essential. We could expect a similar mechanism in the uropathogenic strain UTI89 regarding *pncB* (nicotinate phosphoribosyltransferase), a gene involved in the biosynthesis of NAD. UTI89 is known to require nicotinamide for growth because of a mutation in *nadB*^48,49^. This mutation could also make *pncB* essential by epistatic interaction.

### Mobile genetic elements can trigger the essentiality of core genes

In order to better understand why some genes become essential under certain genetic backgrounds, we then set up a pipeline to isolate mutants that suppress strain-specific essentiality. Briefly, we implemented a two-step selection process using CRISPRi vectors with different resistance markers in order to ensure that the selected mutants did not just inactivate the CRISPRi system (**Fig. 6a, see Methods**). We then sequenced the genome of a few mutants after verifying the absence of mutation in the chromosomal target of the sgRNA which may abrogate repression. In this way, we isolated mutants of the environmental strain E101 that are able to grow with an sgRNA targeting the AAA+ protease *clpP* which is essential in E101 and TA054 (**Fig. 4**). Whole-genome sequencing of 4 suppressor mutants revealed that a prophage also present in TA054 was excised in 3/4 clones and that a hypothetical protein (E101_02645, accession: WP_001179380.1) from the same prophage had a non-synonymous (T46S) mutation in the last clone (**Fig. 6b**). This protein has no predicted domain but seems to be the toxic component of a putative toxin-antitoxin (TA) module with the downstream protein E101_02644 (accession: WP_000481765.1) (these proteins respectively share 33% and 42% of identity to the two components of a putative TA module from *Vibrio mimicus*^50^). Targeting E101_02645 with dCas9 partially rescued the toxicity associated with *clpP* repression in E101 (**Fig. 6c**), while heterologous expression of this system made *clpP* essential in K-12 unless the T46S mutant was expressed (**Fig. 6d**). This confirms that the essentiality of *clpP* in E101 (and probably in TA054) is caused by this putative TA module whose biological role remains to be elucidated.

**Figure 6.**
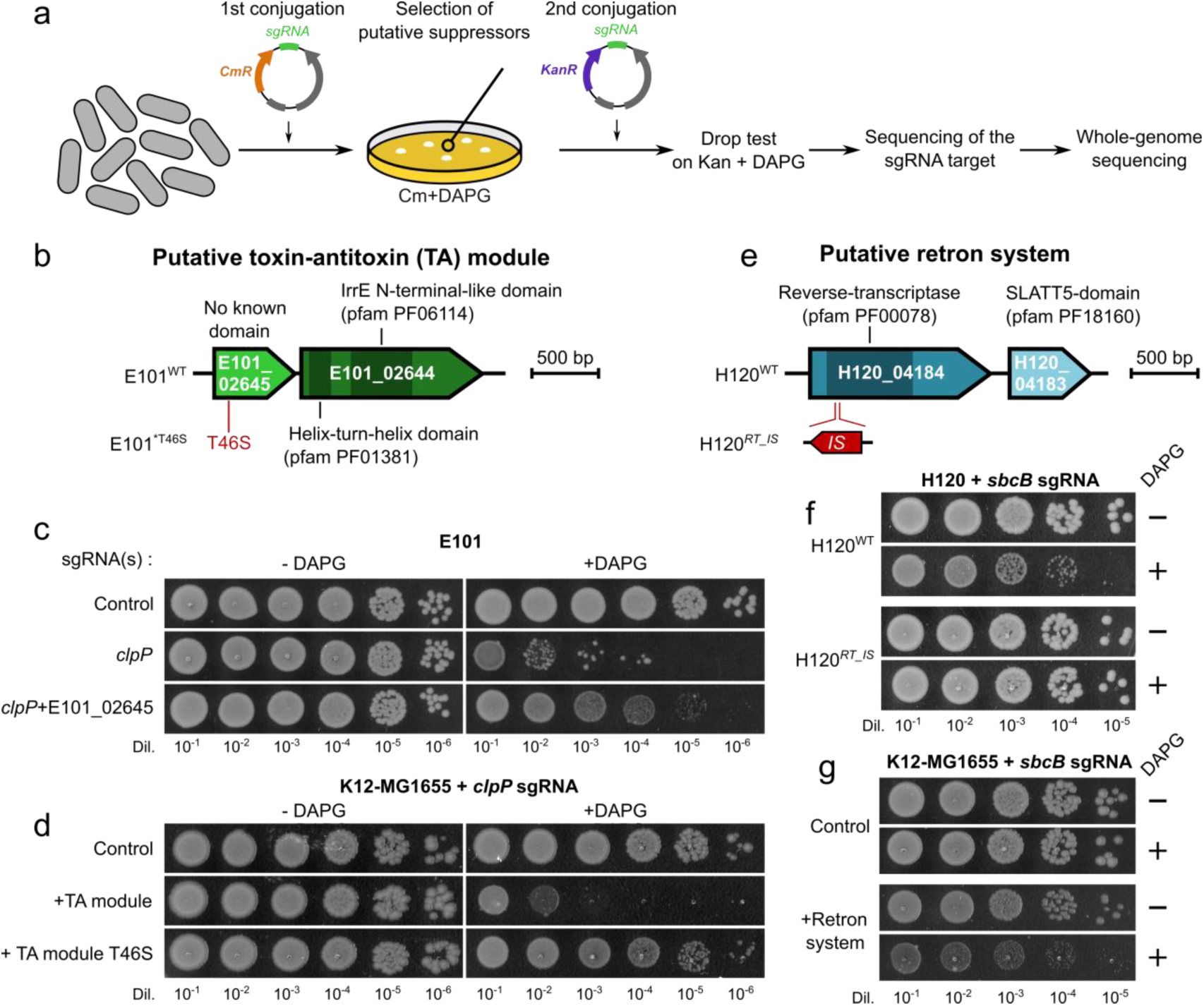
Mapping the genetic determinants of strain-specific gene essentiality. **a**, Suppressor mutants for strain-specific essential genes were isolated by conjugating pFR56 expressing the corresponding sgRNA and selecting transconjugants with chloramphenicol and DAPG to express dCas9. Growing clones were selected and conjugated with a second plasmid with the same sgRNA but with a kanamycin resistance cassette. The resistance to knockdown was verified with a drop assay on kanamycin with DAPG. This ensures that selected clones do not simply carry a mutation in the CRISPRi components. The chromosomal target of the sgRNA was then Sanger-sequenced to verify the absence of mutations abrogating dCas9 binding, before whole-genome sequencing. **b**, Suppressors of *clpP* essentiality in E101 involved a putative toxin-antitoxin module comprising a putative toxin (E101_02645) and its antidote (E101_02644). **c**, Drop assay showing that the toxicity associated with *clpP* repression in E101 is alleviated when E101_02645 is repressed simultaneously. **d**, The putative TA module from E101 WT or from the *clpP* suppressor (harboring the T46S mutation in E101_02645) was transferred to K-12 MG1655. *clpP* knockdown induces a fitness defect with the wild-type TA module but not with the T46S variant. **e**, Suppressors of *sbcB* essentiality in H120 involved a putative retron system encoded by H120_04184 and H120_04183. A suppressor mutant (H120^*RT_IS*^) acquired an insertion element (IS) in the putative reverse-transcriptase gene. **f**, Drop assays showing that the mutant is indeed resistant to *sbcB* knockdown. **g**, Once transferred in K12-MG1655, this system induces a fitness defect when *sbcB* is repressed.

Using the same strategy, we isolated mutants of the uropathogenic strain H120 that can grow in the presence of an sgRNA targeting *lon* (Lon protease) or *sbcB* (exodeoxyribonuclease I). While *sbcB* is essential in H120 and E101, *lon* is essential in H120 only (**Fig. 4**). Similarly to *clpP* in E101, suppressors of *lon* essentiality in H120 had a prophage excised. Two suppressors of *sbcB* essentiality had an insertion element in a protein from the same prophage (H120_04184, accession: WP_000344414.1) that has a reverse-transcriptase domain (pfam 00078) (**Fig. 6e-f**). This gene is believed to be part of a retron system together with the downstream protein (H120_04183, accession: WP_001352776.1) which contains transmembrane helices and a SLATT domain (pfam 18160) that is predicted to function as a pore-forming effector initiating cell suicide^51^. Heterologous expression of this system in K-12 induced a fitness defect when *sbcB* was repressed (**Fig. 6g**). Interestingly, a similar system is present in E101 in the same prophage but associated with a different effector protein. We can hypothesize that this system is also responsible for the essentiality of *sbcB* in E101. The mechanism responsible for the observed fitness defect and the interaction between SbcB and this system remain to be elucidated.

Finally, we identified suppressors of *rnhA* (RNAse HI) essentiality in the ExPEC strain JJ1886. RNAse HI cleaves DNA-RNA duplexes and is involved in DNA replication and repair. We found a suppressor of *rnhA* essentiality in JJ1886 that carries a mutation in the *mfd* gene (transcription-repair coupling factor) and an insertion element in the *rnlA* toxin. This toxin is part of the *rnlAB* TA system and shares 72% of identity with the *E. coli* K-12 RnlA toxin where it is carried by the CP4-57 prophage. In K-12, RnlA has an endoribonuclease activity that is activated upon phage infection to trigger the degradation of phage T4 mRNAs^52^. Strikingly in our strain panel, this TA system is only present in JJ1886 and in K-12 MG1655 (with 72% of identity) which are the only strains where *rnhA* is essential (**Fig. 4**). A recent study in K-12 showed that RNAse HI is required not only to activate the RnlA toxin during phage infection but also to recruit the antitoxin RnlB in uninfected cells to repress RnlA toxicity^53^. The toxicity of *rnhA* knockdown in JJ1886 and K-12 MG1655 is therefore probably caused by the activation of RnlA. The other mutation in *mfd* may also be involved in the suppression of *rnhA* essentiality. Taken together, these findings show that the acquisition of genes by HGT can modify the essentiality of core genes.

## Discussion

We developed a new CRISPRi platform that is compatible with most *E. coli* isolates and with closely-related species of Enterobacteria such as *E. albertii, E. fergusonii, K. pneumoniae* and *C. freundii*. One of the main advantages of CRISPRi over previous techniques is the possibility to design custom sgRNA libraries targeting a subset of genes of interest. CRISPRi screens offer the possibility to easily assess the effect of the same perturbations in different genetic backgrounds. We exploited this feature to design the EcoCG library, a compact library targeting *E. coli* genes that are present in >90% of sequenced isolates. This library is smaller than typical transposon libraries by an order of magnitude. This facilitates multiplexing and substantially decreases sequencing costs. In fact, all screening results presented in this study were obtained from a single Illumina NextSeq 500 run, representing a cost of less than 20€ per sample. We used this library in a panel of 18 isolates from various origins and pathogenicity levels to investigate how the genetic diversity of *E. coli* at the strain level influences the essentiality of conserved genes. Our screens highlighted that the essential character of a gene depends on the genetic background and growth condition, extending recent results in bacteria and in yeast^8–14,18^. The analysis of strains representative of *E. coli* genetic diversity was essential since we found that many phenotypes associated with CRISPRi knockdowns in the model strain K-12 do not translate to other strains.

While screening experiments typically focus on laboratory media, the use of a diverse set of growth conditions is crucial to unveil conditional essential genes. Importantly, many genes that are dispensable in rich laboratory medium are required for *in vivo* growth and conversely^25–29^. Growth media that mimic *in vivo* conditions, such as the GMM^54^ used here, are a valuable alternative to *in vivo* screening experiments that remain difficult to implement on a large scale with a high number of isolates. A recent study in *P. aeruginosa* showed that combining several isolates and growth conditions can highlight essential genes that are specific from infection types, a useful insight for the design of improved therapies^11^. Here, the sets of “variably essential” genes were different across growth conditions and were often associated with a broad range of fitness defects. Instead of a binary trait, gene essentiality should be considered as an extreme fitness defect within a range of continuous values associated with gene disruption. Variations in environmental conditions and genetic backgrounds may modulate the fitness defect of a mutant, or even make the mutation neutral. Our screen, by effectively looking at fitness effects, is a powerful tool to query how natural selection is impacted by these two factors.

We used this data to define the core-essential and the pan-essential genome of *E. coli* comprising genes that are essential in all strains or in at least one strain respectively. Our screens highlighted that when adding new strains, more core genes are susceptible to be found as essential in at least one isolate. As revealed in Figure 2, this trend does not reach a plateau with 18 strains and additional screens of other strains are likely to reveal novel essential genes. This suggests that we are only scratching the surface of the existing differences in gene essentiality at the species level. Note that here we do not report strain-specific essential genes present in the pangenome such as antitoxins and phage repressors, as by design our sgRNA library only targets core genes. We thus underestimate the size of the pan-essential genome.

Differences in gene essentiality correlated poorly with the phylogenetic distance between strains, suggesting that they are not caused by simple processes of vertical evolution and genetic drift. This is in sharp contrast with the strong effect of vertical evolution in the observed variations of gene expression level in our experimental conditions. The investigation of specific differences in gene essentiality revealed three main cases: (i) a few genes such as *kdsB* and *dut* are essential in most strains but dispensable in one or a few strains; (ii) conversely, some genes such as *lon, clpP, sbcB* and *rnhA* are dispensable in most strains but are essential in one or a few strains; (iii) finally some genes such as *rpoE* show a wide range of fitness defects when repressed. In most of the cases that we investigated, gene loss and accretion by horizontal transfer had a key role in providing genetic backgrounds that explain the observed differences.

A gene carrying out an essential function can be dispensable if functional redundancy is provided by another gene. Acquisition or loss of genetic redundancy can thus lead to variations in essentiality. For instance, the inactivation of the core gene *ilvG* in K-12 makes the core operon *ilvHI* essential in minimal medium. This specific event might be linked to the laboratory evolution of strain K-12, as a change in essentiality due to the loss of redundancy between a pair of core genes is likely a rare event. We hypothesized that changes of functional redundancy due to genes of the accessory genome would be more frequent. Homologs of essential genes are indeed common in the accessory genomes of the strains assayed in this work, but we found few cases where this led to changes in essentiality. There are probably several causes for the rarity of effective compensation of an essential gene by a homolog: in several cases, homologs were insufficiently expressed to compensate the loss of the essential gene. In addition, many homologs actually have different molecular functions and are not exchangeable, precluding the compensation of gene loss^44,55^. The search for functionally-redundant genes is also made harder by the fact that compensation can be provided by non-homologous genes. For instance, we describe here how a MazG-like dUTPase (WP_085453089.1) can make *dut* nonessential. In the few cases detailed here, the accessory genes compensating the loss of an essential gene were acquired in mobile genetic elements. The residence time of mobile elements in the genome is usually short. As a result, the permanent replacement of an essential gene by a functionally-redundant gene could be thought as an unlikely contingency. Yet, cases of non-orthologous gene displacement, i.e. replacement of a key gene by a distant homolog or an analog, are frequent at the macro-evolutionary scale^56^. The functional redundancy provided by incoming mobile elements that we describe here might thus be a first step towards actual gene replacement on a longer evolutionary timescale.

Beyond gain and loss of functional redundancy, the essentiality of a function itself might change due to genetic differences between strains. For instance, a mutation in the ribosome release factor 2 encoded by *prfB* in K-12 MG1655 leads to dramatically impaired translation termination in the absence of a modification of the 50S ribosome subunit carried out by the product of *rluD*, making this gene essential^47^. Previous studies also showed that the loss of some essential genes can be compensated by the overexpression of genes carrying a different function^57,58^. We found here that gene loss and accretion can also modify the essentiality of functions carried in the core genome. The isolation of suppressor mutants of strain-specific essential genes led us to the identification of genes encoded on mobile elements that make genes such as *clpP, sbcB*, or *rnhA* essential. Overall, our data shows the ability of some core genes to become essential on a recurrent basis after HGT events, a phenomenon that could favor the evolutionary conservation of these genes despite their dispensability in most conditions. Since this phenomenon is likely linked to the high rate of HGT in *E. coli*, we could expect extensive differences in gene essentiality in any species with an open pan-genome.

## Methods

### Bacterial cultivation

Unless stated otherwise, lysogeny broth (LB) broth was used as a liquid medium and in LB + 1.5 % agar as a solid medium. Kanamycin (Kan) was used at 50 μg / mL, chloramphenicol (Cm) was used at 20 µg/mL, erythromycin (Erm) was used at 200 µg/mL and carbenicellin was used at 100 μg / mL. The composition of media used for screening is described in **Supplementary Table 2**. *E. coli* K-12 MG1655 was used for cloning and MFDpir was used for plasmid transfer by conjugation^42^.

### Plasmid construction

The dCas9-sgRNA plasmid expression system, pFR56, was derived from plasmid pJF1, a gift from Eligo Bioscience, harbouring a constitutively expressed sgRNA and cas9 under the control of a DAPG-inducible PhlF promoter. This plasmid was recoded to avoid restriction sites in order to ensure plasmid stability in a maximum of *E. coli* strains^38^. We further modified this plasmid to inactivate Cas9 into dCas9 and add the RP4 origin of transfer. Novel sgRNAs can be cloned on pFR56 using Golden Gate assembly^59^ with BsaI restriction sites. The expression level of dCas9 on pFR56 was optimized to avoid the previously reported toxicity effect, known as the bad-seed effect^33^. Briefly, we used the RBS calculator^60^ to randomize 4 positions of the dCas9 RBS and cloned the resulting library on the plasmid harboring an sgRNA with a bad-seed sequence (5’-TTGTATCAAACCATCACCCA-3’) using the Gibson assembly method^61^. Candidate clones that grew normally in the presence of dCas9 induction were selected. In order to select clones that retain a sufficient dCas9 expression level for efficient repression of target genes, an sgRNA targeting the essential gene *rpsL* cloned onto the psgRNAcos vector (Addgene accession 114005) was introduced in the selected candidates. We discarded clones that were not killed in the presence of dCas9 induction. Finally the sgRNA was modified to include a *ccdB* counter-selection cassette in between two BsaI restriction sites^62^. This ensures the selection of clones in which a guide was successfully added to the plasmid during library cloning.

### Library design

We retrieved all *E. coli* complete genomes from GenBank Refseq (available in February 2018). We estimated genome similarity calculating the pairwise Mash distance (M) between all genomes using Mash v.2.0^63^. Importantly, the correlation between the Mash distances (M) and Average Nucleotide Identity (ANI) in the range of 90-100% has been shown to be very strong, with M ≈ 1-(ANI/100). All the resulting Mash distances between *E. coli* genomes are well below 0.05, in agreement with the assumption that they all belong to the same species. We just removed some genomes that were too similar (MASH distance < 0.0001). In this case, we picked the one present for a longer period of time in the databases. This resulted in a dataset of 370 completely assembled genomes for comparison^22^. Pan-genomes are the full complement of genes in the species (or dataset, or phylogroup) and were built by clustering homologous proteins into families. We determined the lists of putative homologs between pairs of genomes with MMseqs2 v.3.0^64^ by keeping only hits with at least 80% identity and an alignment covering at least 80% of both proteins. Homologous proteins were then clustered by single-linkage^65^. From the resulting pangenome, we selected 3380 proteins present in more than 333/370 genomes (90%) in up to 4 copies per genome.

For each gene and strain, all possible sgRNAs were listed by selecting the 20 NGG-proximal nucleotides on the coding strand. In order to avoid sgRNAs targeting regions with single-nucleotide variants, a first pre-selection step was performed for each gene in order to select up to 12 sgRNAs based on the number of targeted strains: (i) sgRNAs targeting the highest number of strains (N_max_) were first selected; (ii) if less than 12 guides were obtained, sgRNAs targeting N_max_-1 strains were selected, then N_max_-2 strains, etc until 90% x N_max_ strains; (iii) if less than 3 sgRNAs were selected after this process (possibly due to high rates of variants), the 3 sgRNAs with the highest number of targeted strains were selected; (iv) in order to select sgRNAs targeting the strains that may have been missed, we then selected the strains targeted by less than 3 sgRNAs and performed a similar selection procedure: sgRNAs targeting the maximum number of missed strains (N_max_missed_) were selected, followed by sgRNAs targeting N_max_missed_-1 strains, etc, until 80% x N_max_missed_ strains. Finally, sgRNAs targeting less than 30 strains (∼8%) were discarded.

After the preselection process, a penalty score was calculated from each sgRNA in order to select the best 3 sgRNAs targeting each gene. This score takes into account, (i) off-target effects, (ii) predicted efficiency, (iii) number of targeted strains.

i. For each sgRNA, we calculated the fraction of strains having another 11-nt match on the coding strand of a gene and the fraction of strains having a 9-nt match on any strand in a promoter (loosely defined as 100 nt before gene start). The 1^st^ score was calculated as the sum of these fractions.
ii. We used a recent model^39^ which predicts the repression efficiency of sgRNAs based on fitness data obtained in a previous CRISPRi screen^33^. For each gene, the predicted sgRNA activity was normalized from 0 (highest activity) to 1 (lowest activity) and was then used as a 2^nd^ score.
iii. The number of targeted strains (with a full-length match) was reported for each sgRNA. For each gene, this number was normalized from 0 (sgRNA targeting the most strains) to 1 (virtually no strain targeted) and was then used as a 3^rd^ score.

For each gene, all preselected sgRNAs were attributed a global penalty score by summing the 3 scores described above. A strong penalty was applied to guides carrying a 5-nt seed sequence among the 10 strongest bad-seed sequences identified by Cui *et al*. (2018)^33^ (AGGAA, TAGGA, ACCCA, TTGGA, TATAG, GAGGC, AAAGG, GGGAT, TAGAC, GTCCT), so that they were only selected as a last resort. For each gene, sgRNAs were ranked by increasing global penalty score and the 3 best sgRNAs were selected (if available). If one of these 3 sgRNAs targeted less than 350 strains (95%), a 4^th^ sgRNA was added. This process resulted in a library of 11,188 sgRNAs targeting conserved protein-coding genes.

We also designed sgRNAs targeting rRNAs, tRNAs and ncRNAs. Since rRNAs are highly conserved between all strains, it is very simple to select sgRNAs targeting all strains. However, it is complicated to assess their potential off-target activity due to their presence in many copies. We therefore selected all 109 sgRNAs targeting all strains. Similarly, homologous tRNAs have very similar nucleotide sequences, which makes it difficult to assess the off-target activity of each sgRNA. We therefore selected 131 sgRNAs targeting > 90% of strains (> 333 strains). ncRNAs are very diverse and their annotation can substantially differ between genomes. All annotated ncRNAs in all genomes were first listed. Then, ncRNAs which sequence is present in > 100 genomes were kept. For each of these ncRNAs, all possible sgRNAs were listed and assessed for off-target activity. For each ncRNA, we determined the off-target size *s* for which at least 3 guides had no off-target activity (or in < 5% of strains). We then selected all the guides having an off-target of size *s* in < 5% of strains.

Finally, we generated 20 non-targeting control sgRNAs. These guides should not have any activity nor off-target in any strain. To ensure a minimal off-target activity, we first generated all possible random 8-mers and kept those which never occur next to a PAM in a subset of 20 strains. We then generated 20 sgRNAs whose 8 last 3’ bases were randomly chosen from this subset, and whose 12 first 5’ bases were random. We verified that each control sgRNA does not have an off-target in more than 1% of strains.

### Library construction

The resulting library of 11,629 sgRNAs was generated as single-stranded DNA through on-chip oligo synthesis (CustomArray). Pooled oligo extension was performed with KAPA HiFi DNA polymerase (Roche) with primer FR222. The library was then amplified by PCR (KAPA HiFi polymerase; 95°C - 3’; 6 cycles 98°C – 20”, 60°C – 15”, 72°C – 20’’; 72°C – 10’) with primers FR221 and FR222 and purified by gel extraction. The pFR56-ccdB vector was digested with BsaI (New England Biolabs) and gel purified. The plasmid library was then assembled using the Gibson method^61^.

During transformation, the initial absence of repressor proteins in the cell can result in a transient dCas9 expression which can introduce a bias in the library. To avoid this, we built a library cloning strain, FR-E03, by integrating a constitutively-expressed PhlF repressor gene in the chromosome of MG1655. Briefly, a phlF expression cassette was cloned onto the pOSIP backbone^66^ and integrated into HK022 *attP* site. The pOSIP backbone was then excised using the pE-FLP plasmid which was cured by serial restreaks. For transformation of the library, FR-E03 cells were grown in LB (200 mL) to OD ∼ 1, washed 3 times in ice-cold pure water and resuspended in 250 µL ice-cold water. Ten electroporations were performed with 20 µL cells and 0.5 µL of dialyzed Gibson assembly product and pooled. After 1h at 37°C, cells were plated on 10 large LB-Cm plates (12×12 cm) and incubated overnight at room temperature. The next day, each plate was washed twice with 5 mL LB-Cm and pooled. Plasmids were extracted by miniprep (Mancherey-Nagel) before further transformation into the conjugation strain MFDpir. This *pir*+ strain is auxotrophic to diaminopimelic acid (DAP) and contains the RP4 conjugation machinery^42^. We attempted to integrate the same construction as in FR-E03 in the conjugation strain MFDpir but this was unsuccessful. Instead, we used pFR58, a low-copy pSC101 Kan^R^ plasmid with the same PhlF expression system. We confirmed that pFR58 cannot be mobilized during conjugation since it does not contain the RP4 transfer machinery. The library was therefore electroporated into MFDpir+pFR58 as described above. Transformants were selected on LB agar supplemented with Cm and 300 µM DAP and pooled before conjugation.

### Strain selection

Starting from a collection of 92 *E. coli* natural isolates encompassing the phylogenetic diversity of the species and originating from various habitats in diverse conditions (environment, birds, non-human mammals and humans; gut commensalism, intestinal and extra-intestinal infections), we performed growth curves to identify natural resistance to chloramphenicol which is the selection marker used in pFR56. Briefly, overnight cultures were diluted 100-fold in LB or in LB-Cm and OD600 was measured every 10 min for 8 h at 37°C with shaking on a Tecan Infinite M200Pro. Successful growth in Cm was observed in seven strains which were discarded. dCas9-mediated repression was then tested by conjugating pFR56 bearing an sgRNA targeting the essential gene *rpsL* into each strain. Plating on LB-Cm + 50 µM DAPG induced strong killing in all strains, suggesting that dCas9-mediated repression is functional in all strains. From the remaining isolates, we selected a panel of 18 strains including K-12 MG1655 from diverse origin and pathogenicity spanning most common *E. coli* phylogroups (A, B1, B2, D, E and F). Phylogroups were verified by quadruplex PCR with the Clermont method^67^. Strains selected for screening are listed in **Supplementary Table 1**.

### Bacterial conjugation

MFDpir cells carrying the plasmid library were grown to OD∼1 in LB-Cm supplemented with 300 µM DAP. Cells were then washed (2,000g – 10’) to remove traces of Cm. Recipient strains were grown to stationary phase. Donor and recipients cells were mixed 1:1 (v/v, 0.1 to 1mL), pelleted (2,000g – 10 min), resuspended in 10-100 µL LB + 300 µM DAP, pipetted onto a LB + 300 µM DAP plate and incubated at 37°C for 2 h. As a negative control, donor and recipient strains were plated on a LB-Cm plate. For conjugation of individual sgRNAs, cells were then restreaked on LB-Cm to select individual 6transconjugants. For conjugation of the EcoCG library, cells were collected, resuspended in 1 mL of LB-Cm and plated on a large LB-Cm plate (12 × 12 cm) followed by overnight incubation at room 6temperature. Ten-fold dilutions were also plated for CFU counting. We obtained >10^7^ clones for each of the 18 strains assayed in this study, ensuring a > 1000-fold coverage. After overnight incubation at room temperature, plates containing nascent colonies were washed twice in 5 mL LB-Cm and stored at −80°C with 7.5% DMSO.

### Screen design

Strains conjugated with the library were arrayed by mixing 150 µL of the −80°C stock with 1350 µL LB-Cm in duplicates on a 96-deepwell plate (Masterblock 96 well, 2ml, V-bottom plates by Greiner Bio-one). The plate was incubated overnight at 37°C in a Thermomixer (Eppendorf) with shaking (700 rpm). The next day, cultures were washed 1:1 in M9 medium before inoculation of 15 µL into 1485 µL of either LB, M9-glucose or GMM, supplemented with 50 µM DAPG without antibiotic selection. The remaining cultures were harvested and plasmids were extracted by miniprep to obtain reference samples for each strain. All screens were then performed in a Thermomixer at 37°C with shaking (700 rpm). Screens in LB and M9-glucose were performed in aerobic condition while screens in GMM were performed in an anaerobic chamber (80% N_2_, 10% CO_2_ and 10% H_2_). In all three cases, 3 passages were performed by diluting 15 µL of cells into 1485 µL of DAPG-containing fresh medium (1:100 6dilution) in the same conditions, every 3.5 h for LB and every 12 h for M9-glucose and GMM. This represents a total of ∼20 generations (log2(100^3^)≈19.9). Plasmids were finally extracted with a 96-well miniprep kit (Macherey-Nagel) to obtain the final distribution of the library for each strain and medium.

### Illumina sample preparation and sequencing

Library sequencing was performed as previously described^33,34^. Briefly, primers listed in **Supplementary Table 6** were used to perform two consecutive PCR reactions with KAPA HiFi polymerase (Roche). Starting from 100 ng of library plasmid, the first PCR (95°C – 3 min; 9 cycles [98°C – 20 s; 60°C – 15 s; 72°C – 20 s]; 72°C – 10 min) is performed in a 30-µL reaction with 8.6 pmol of each primer. For the second PCR, a 20-µL mix containing 100 pmol of primers is added to the first PCR and the resulting 50-µL reaction is incubated (95°C – 3 min; 9 cycles [98°C – 20 s; 66°C – 15 s; 72°C – 20 s]; 72°C – 10 min) to add the 2nd index and the flow cell attachment sequences. The resulting 354 bp-PCR DNA fragments were gel extracted. Samples were pooled (150 ng of each reference samples and 100 ng of other samples) and the final library concentration was determined by qPCR (KAPA Library Quantification Kit, Roche). Sequencing was performed on a NextSeq 500 benchtop sequencer (Illumina) using a custom protocol as previously described^34^. We obtained an average of 3 million reads per sample, representing an average coverage of ∼ 260X.

### Data analysis

Index sequences were used to de-multiplex the data into individual samples with a custom Python script. Reproducibility between experimental duplicates was very high (median Pearson’s r = 0.988) except for a replicate of strain ROAR 8 in LB that had very low read counts. This sample was discarded, while biological replicates from other strains were pooled into a single sample per strain for subsequent analyses. sgRNAs with less than 20 reads in the initial time point of a given strain were discarded in the corresponding strain (1.9% of the library on average). Samples were then normalized by sample size. The log2FC value was calculated for each guide *g* and strain *s* as follows (*s_initial* and *s_final* represent the normalized reads counts of strain *s* in the initial and final time point respectively):

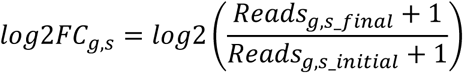

We mapped the EcoCG library to each genome to identify sgRNAs which do not have a full-length match (e.g. because of single-nucleotide variants), and their log2FC value was set to NaN in the corresponding strain in order to avoid false negatives. Finally, the median log2FC were centered on the median log2FC of 20 control non-targeting sgRNAs, and the resulting values were used as a gene scores. We selected genes as “variably essential” in **Supplementary Fig. 5** when repression induced a fitness defect in at least one strain (gene score < −3) and no fitness defect in at least one strain (gene score > −1). For the heatmaps drawn in **Fig. 4**, we used a more stringent threshold: for each gene, we calculated the minimum and maximum gene scores across strains after excluding those that had more than 50% of sgRNAs with missing values. We then kept genes whose minimal gene score was lower than −5 and whose maximal gene score was greater than −1 across all strains.

### Comparative genomics

The genomes of the 18 strains used for screening were reannotated with Prokka 1.14.2 using default settings^68^. Proteins from these 18 strains were clustered using MMseqs2 v.3.0 with default parameters^64^. The resulting clusters were used to generate the core and pan-genome shown in **Fig. 2** using up to 250 permutations of sets of strains. To obtain pairwise phylogenetic distances between strains, we generated a core genome alignment with Parsnp^69^ which was used to build a phylogenetic tree with FastTree2^70^. To evaluate the presence of homologs of essential genes, we clustered proteins from all 18 strains with a 40%-identity threshold with MMseqs2 v.3.0^64^ (--min-seq-id 0.4) to obtain groups of sequence homologs. We then selected clusters containing essential proteins from K12-BW25113 in LB^17^. To do so, we clustered proteins from BW25113 and MG1655 (--min-seq-id 0.95) to obtain a correspondence table between the names of BW25113 essential genes and MG1655 locus tags. We finally selected protein clusters containing at least one sequence per strain with at least one strain having more than one sequence.

We used sequences searches on the web interface of InterPro^71^ and pfam^72^ databases to look for known domains in protein candidates. Structural predictions were performed with Phyre2^73^.

### Screen results validation

#### sgRNA cloning

Individual sgRNAs listed in **Supplementary Table 7** were cloned into pFR56 using Golden Gate assembly^59^. All constructions were validated by Sanger sequencing. Cloning was performed in MG1655 or MFDpir before transfer to the appropriate strains by conjugation.

#### Gene deletions

Genes *dut* and *rpoE* were deleted from strain TA447 using the λ-red system recombination system as described previously^34^ using primers listed in **Supplementary Table 6**. We performed whole-genome sequencing (WGS) to verify the absence of compensatory mutations. We used breseq (v. 0.33.2) for variant calling^74^.

#### Growth curves

For growth curves, an overnight culture was washed in the appropriate medium to avoid nutrient carryover and diluted 1000-fold. Growth was monitored in triplicates by measuring optical density at 600 nm on an Infinite M200Pro (Tecan) at 37°C with shaking.

#### RT-qPCR

Overnight cultures of S88 carrying pFR56 or pFR56.27 (i.e. pFR56 with *ybaQ* sgRNA) in LB + 692 Cm were diluted 1000-fold in 2 mL of LB + Cm + 50 µM DAPG. An overnight culture of TA447 was 6diluted 1000-fold in 2 mL of LB and 100-fold in 2 mL of GMM to account for the slower growth rate in GMM. After 3h at 37°C, RNA were extracted using Trizol. RNA samples were treated with DNAse (Roche) and reverse-transcribed into cDNA using the Transcriptor First Strand cDNA Synthesis Kit (Roche). qPCR was performed in two technical replicates with the FastStart Essential DNA Green master mix (Roche) on a LightCycler 96 (Roche). Relative gene expression was computed using the ΔΔCq method after normalization by 5S rRNA (*rrsA*). qPCR primers are listed in **Supplementary Table 8**.

#### Isolation of suppressor mutants

To isolate suppressors mutants, we conjugated pFR56 harboring the corresponding guide into the appropriate strain and we selected clones that grew robustly with 50 µM DAPG. In order to avoid selecting mutations on the plasmid that inactivate the CRISPRi system, we conjugated a second plasmid (pFR59) identical to pFR56, but carrying a kanamycin resistance cassette instead of the chloramphenicol resistance cassette. The resistance to repression was verified by plating serial dilutions of the transconjugants on LB agar plates with Kan ± 50 µM DAPG. In the case of JJ1886, this strain is naturally resistant to kanamycin. We therefore built a third plasmid (pFR72) with a gentamycin resistance cassette and used it for the first selection step together with pFR56 in the second selection step. Finally, to avoid selecting clones that acquired mutations in the chromosomal sgRNA target, we performed Sanger sequencing on the genomic region flanking the sgRNA binding site and discarded clones with mutations in the target. Genomic DNA was extracted from selected clones as well as in the parental clone using the Wizard Genomic DNA Purification Kit (Promega). NGS was performed using Nextera XT DNA Library Preparation kit and the NextSeq 500 sequencing systems (Illumina) at the Mutualized Platform for Microbiology (P2M) at Institut Pasteur. Mutations were identified by mapping raw reads to the appropriate genome using breseq v. 0.33.2^74^. Among the genomes we sequenced, APEC O1 had a previously unreported plasmid. The new genome was deposited on the European Nucleotide Archive (ENA) under the accession GCA_902880315. We also found that the previously reported genome sequence of H120 (GCF_000190855.1) had a high number of sequencing errors introducing frameshifts and premature stop codons. We resequenced this strain to correct these errors and deposited the resulting corrected genome sequence on the ENA with the accession GCA_902876715.

### RNA-seq

Overnight cultures were diluted 100-fold in 1 mL of LB in a 96-deepwell plate (Masterblock 96 well, 2ml, V-bottom plates by Greiner Bio-one). After 2.5 h at 37°C, cultures were diluted to OD ∼ 0.02 in 1.4 mL of LB and were further grown for 2 h at 37°C on a Thermomixer (Eppendorf) with shaking (700 rpm). Each culture was then transferred to a 2-mL Eppendorf containing 170 µL of stop solution (5% acid phenol in ethanol) and cooled down for 10 seconds in a bath of dry ice and ethanol. Cells were 7harvested by centrifugation at 0°C (1 min – 16,000 g) and the pellets were frozen at −80°C. For RNA extraction, pellets were thawed on ice, resuspended in 200 µL of pre-warmed lysozyme solution and incubated for 3 min at 37°C before addition of 1 mL of Trizol. Samples were vigorously vortexed and incubated at room temperature for 5 min followed by addition of 200 µL of chloroform. After vigorous vortexing, samples were incubated for 5 min at room temperature and centrifuged (10 min – 12,000 g) to separate phases. The upper aqueous phase was collected and RNA was precipitated by addition of 500 µL of isopropanol. Samples were incubated for 10 min at room temperature before centrifugation (10 min – 12,000 g). Pellets were washed with 1 mL of 75% ethanol and centrifuged (5 min – 7,500 g). The pellets were finally air-dried and resuspended in 50 µL of pure water. RNA samples were DNAse-treated using TURBO DNA-free Kit (Thermo Fisher Scientific) and sample quality was assessed on an Agilent Bioanalyzer 2100. Samples were prepared for sequencing using the TruSeq^®^ Stranded Total RNA Kit (Illumina) and sequenced on a NextSeq 500 benchtop sequencer (Illumina). Raw reads were aligned on each genome using Bowtie2 v2.3.4.3^75^. Alignment files were converted with Samtools v1.9^76^ and read counts for each gene were obtained using HTseq v0.9.1^77^. Raw read counts were normalized by sample size and by gene length to obtain reads-per-kilobase-per-million (RPKM). The log2-transformed median RPKM value of each across biological replicates was used as a measure of gene expression in each strain.

### Data availability

Raw sequencing reads from CRISPRi screens are available at the European Nucleotide Archive under the accession PRJEB37847. Raw reads from RNA-seq experiments were deposited on ArrayExpress with the accession E-MTAB-9036. Processed data is available in Supplementary Tables.

## Acknowledgement

We thank Bruno Dupuy for sharing the use of the anaerobic chamber, as well as Olivier Tenaillon, Pierre-Alexandre Kaminsky, Guennadi Sezonov, Antoine Danchin and Alicia Calvo-Villamañan for useful discussions. We thank the P2M platform (Institut Pasteur, Paris, France) for genome sequencing and the Biomics platform (Institut Pasteur, Paris, France) for preparing RNA samples for sequencing. This work was supported by the European Research Council (ERC) under the Europe Union’s Horizon 2020 research and innovation program (grant agreement No [677823]), by the French Government’s Investissement d’Avenir program and by Laboratoire d’Excellence ‘Integrative Biology of Emerging Infectious Diseases’ (ANR-10-LABX-62-IBEID), F.R. is supported by a doctoral scholarship from Ecole Normale Supérieure. E.D. was partially supported by the “Fondation pour la Recherche Médicale” (Equipe FRM 2016, grant number DEQ20161136698). E.R. was partially supported by the “Fondation pour la Recherche Médicale” (Equipe FRM EQU201903007835).

## Author contributions

F.R. and D.B. designed the project. E.R. performed bioinformatic computation of the *E. coli* pangenome. E.D. and O.C. provided strains and genome sequences. F.R. performed experiments and analyzed data. J.R.F. and F.P.F. participated in the design of pFR56. J.C.C. provided experimental assistance. F.R., E.R. and D.B. wrote the manuscript. D.B. supervised the project.

## Competing interests

The authors declare no competing interests.

## Notes

### Competing Interest Statement

The authors have declared no competing interest.

## References

1. Rancati, G., Moffat, J., Typas, A. & Pavelka, N. Emerging and evolving concepts in gene essentiality. Nat. Rev. Genet. 19, 34–49 (2017).

2. Kohanski, M. A., Dwyer, D. J. & Collins, J. J. How antibiotics kill bacteria: from targets to networks. Nat. Rev. Microbiol. 8, 423–435 (2010).

3. Jarboe, L. R. Improving the success and impact of the metabolic engineering design, build, test, learn cycle by addressing proteins of unknown function. Curr. Opin. Biotechnol. 53, 93–98 (2018).

4. Hutchison, C. A. et al. Design and synthesis of a minimal bacterial genome. Science (80-.). 351, aad6253 (2016).

5. Rocha, E. P. C. & Danchin, A. An Analysis of Determinants of Amino Acids Substitution Rates in Bacterial Proteins. Mol. Biol. Evol. 21, 108–116 (2004).

6. Jordan, I. K., Rogozin, I. B., Wolf, Y. I. & Koonin, E. V. Essential genes are more evolutionarily conserved than are nonessential genes in bacteria. Genome Res. 12, 962–8 (2002).

7. Thomason, M. K. et al. Global transcriptional start site mapping using differential RNA sequencing reveals novel antisense RNAs in Escherichia coli. J. Bacteriol. 197, 18–28 (2015).

8. Turner, K. H., Wessel, A. K., Palmer, G. C., Murray, J. L. & Whiteley, M. Essential genome of Pseudomonas aeruginosa in cystic fibrosis sputum. Proc. Natl. Acad. Sci. U. S. A. 112, 4110–4115 (2015).

9. Le Breton, Y. et al. Essential Genes in the Core Genome of the Human Pathogen Streptococcus pyogenes. Sci. Rep. 5, 9838 (2015).

10. Freed, N. E., Bumann, D. & Silander, O. K. Combining Shigella Tn-seq data with gold-standard E. coli gene deletion data suggests rare transitions between essential and non-essential gene functionality. BMC Microbiol. 16, 203 (2016).

11. Poulsen, B. E. et al. Defining the core essential genome of Pseudomonas aeruginosa. Proc. Natl. Acad. Sci. U. S. A. 116, 10072–10080 (2019).

12. Galardini, M. et al. The impact of the genetic background on gene deletion phenotypes in *Saccharomyces cerevisiae*. Mol. Syst. Biol. 15, (2019).

13. Dowell, R. D. et al. Genotype to phenotype: a complex problem. Science 328, 469 (2010).

14. van Opijnen, T., Dedrick, S. & Bento, J. Strain Dependent Genetic Networks for Antibiotic-Sensitivity in a Bacterial Pathogen with a Large Pan-Genome. PLOS Pathog. 12, e1005869 (2016).

15. Baba, T. et al. Construction of Escherichia coli K-12 in-frame, single-gene knockout mutants: the Keio collection. Mol. Syst. Biol. 2, 2006.0008 (2006).

16. Nichols, R. J. et al. Phenotypic Landscape of a Bacterial Cell. Cell 144, 143–156 (2011).

17. Goodall, E. C. A. et al. The Essential Genome of Escherichia coli K-12. MBio 9, e02096–17 (2018).

18. Price, M. N. et al. Mutant phenotypes for thousands of bacterial genes of unknown function. Nature 557, 503–509 (2018).

19. Wetmore, K. M. et al. Rapid quantification of mutant fitness in diverse bacteria by sequencing randomly bar-coded transposons. MBio 6, e00306–15 (2015).

20. Rasko, D. A. et al. The pangenome structure of Escherichia coli: comparative genomic analysis of E. coli commensal and pathogenic isolates. J. Bacteriol. 190, 6881–93 (2008).

21. Touchon, M. et al. Organised Genome Dynamics in the Escherichia coli Species Results in Highly Diverse Adaptive Paths. PLoS Genet. 5, e1000344 (2009).

22. Touchon, M. et al. Phylogenetic background and habitat drive the genetic diversification of Escherichia coli. bioRxiv 2020.02.12.945709 (2020). doi: 10.1101/2020.02.12.945709

23. Tenaillon, O., Skurnik, D., Picard, B. & Denamur, E. The population genetics of commensal Escherichia coli. Nat. Rev. Microbiol. 8, 207–217 (2010).

24. Hobman, J. L., Penn, C. W. & Pallen, M. J. Laboratory strains of Escherichia coli: model citizens or deceitful delinquents growing old disgracefully? Mol. Microbiol. 64, 881–885 (2007).

25. Subashchandrabose, S., Smith, S. N., Spurbeck, R. R., Kole, M. M. & Mobley, H. L. T. Genome-Wide Detection of Fitness Genes in Uropathogenic Escherichia coli during Systemic Infection. PLOS Pathog. 9, e1003788 (2013).

26. Olson, M. A., Siebach, T. W., Griffitts, J. S., Wilson, E. & Erickson, D. L. Genome-Wide Identification of Fitness Factors in Mastitis-Associated Escherichia coli. Appl. Environ. Microbiol. 84, e02190–17 (2018).

27. Phan, M.-D. et al. The Serum Resistome of a Globally Disseminated Multidrug Resistant Uropathogenic Escherichia coli Clone. PLOS Genet. 9, e1003834 (2013).

28. Goh, K. G. K. et al. Genome-Wide Discovery of Genes Required for Capsule Production by Uropathogenic Escherichia coli. MBio 8, e01558–17 (2017).

29. Shea, A. E. et al. Identification of *Escherichia coli* CFT073 fitness factors during urinary tract infection using an ordered transposon library. Appl. Environ. Microbiol. (2020). doi: 10.1128/AEM.00691-20

30. Martínez-Carranza, E. et al. Variability of Bacterial Essential Genes Among Closely Related Bacteria: The Case of Escherichia coli. Front. Microbiol. 9, 1059 (2018).

31. Qi, L. S. et al. Repurposing CRISPR as an RNA-Guided Platform for Sequence-Specific Control of Gene Expression. Cell 152, 1173–1183 (2013).

32. Bikard, D. et al. Programmable repression and activation of bacterial gene expression using an engineered CRISPR-Cas system. Nucleic Acids Res. 41, 7429–7437 (2013).

33. Cui, L. et al. A CRISPRi screen in E. coli reveals sequence-specific toxicity of dCas9. Nat. Commun. 9, 1912 (2018).

34. Rousset, F. et al. Genome-wide CRISPR-dCas9 screens in E. coli identify essential genes and phage host factors. PLOS Genet. 14, e1007749 (2018).

35. Wang, T. et al. Pooled CRISPR interference screening enables genome-scale functional genomics study in bacteria with superior performance. Nat. Commun. 9, 2475 (2018).

36. Lee, H. H. et al. Functional genomics of the rapidly replicating bacterium Vibrio natriegens by CRISPRi. Nat. Microbiol. 4, 1105–1113 (2019).

37. Yao, L. et al. Pooled CRISPRi screening of the cyanobacterium Synechocystis sp PCC 6803 for enhanced industrial phenotypes. Nat. Commun. 11, 1666 (2020).

38. Decrulle, A., Fernandez Rodriguez, J., Duportet, X. & Bikard, D. OPTIMIZED VECTOR FOR DELIVERY IN MICROBIAL POPULATIONS. (2018).

39. Calvo-Villamañán, A. et al. On-target Activity Predictions Enable Improved CRISPR-dCas9 Screens in Bacteria. Nucleic Acids Res. (2020). doi: 10.1093/nar/gkaa294

40. rchounian, A. & Trchounian, K. Fermentation Revisited: How Do Microorganisms Survive Under Energy-Limited Conditions? Trends in Biochemical Sciences 44, 391–400 (2019).

41. Zhang, J. & Yang, J.-R. Determinants of the rate of protein sequence evolution. Nat. Rev. Genet. 16, 409–420 (2015).

42. Ferrières, L. et al. Silent mischief: bacteriophage Mu insertions contaminate products of Escherichia coli random mutagenesis performed using suicidal transposon delivery plasmids mobilized by broad-host-range RP4 conjugative machinery. J. Bacteriol. 192, 6418–27 (2010).

43. Andersson, D. I. & Hughes, D. Gene Amplification and Adaptive Evolution in Bacteria. Annu. Rev. Genet. 43, 167–195 (2009).

44. ian, W. & Skolnick, J. How well is enzyme function conserved as a function of pairwise sequence identity? J. Mol. Biol. 333, 863–882 (2003).

45. Tye, B.-K. & Lehman, I. R. Excision repair of uracil incorporated in DNA as a result of a defect in dUTPase. J. Mol. Biol. 117, 293–306 (1977).

46. Gao, Y. et al. Systematic discovery of uncharacterized transcription factors in Escherichia coli K-12 MG1655. Nucleic Acids Res. 46, 10682–10696 (2018).

47. Schaub, R. E. & Hayes, C. S. Deletion of the RluD pseudouridine synthase promotes SsrA peptide tagging of ribosomal protein S7. Mol. Microbiol. 79, 331–341 (2011).

48. Li, Z., Bouckaert, J., Deboeck, F., De Greve, H. & Hernalsteens, J.-P. Nicotinamide dependence of uropathogenic Escherichia coli UTI89 and application of nadB as a neutral insertion site. Microbiology 158, 736–745 (2012).

49. Bouvet, O., Bourdelier, E., Glodt, J., Clermont, O. & Denamur, E. Diversity of the auxotrophic requirements in natural isolates of Escherichia coli. Microbiology 163, 891–899 (2017).

50. Luo, P., He, X., Liu, Q. & Hu, C. Developing Universal Genetic Tools for Rapid and Efficient Deletion Mutation in Vibrio Species Based on Suicide T-Vectors Carrying a Novel Counterselectable Marker, vmi480. PLoS One 10, e0144465 (2015).

51. Burroughs, A. M., Zhang, D., Schäffer, D. E., Iyer, L. M. & Aravind, L. Comparative genomic analyses reveal a vast, novel network of nucleotide-centric systems in biological conflicts, immunity and signaling. Nucleic Acids Res. 43, 10633–54 (2015).

52. Otsuka, Y., Ueno, H. & Yonesaki, T. Escherichia coli endoribonucleases involved in cleavage of bacteriophage T4 mRNAs. J. Bacteriol. 185, 983–990 (2003).

53. Naka, K., Qi, D., Yonesaki, T. & Otsuka, Y. RnlB antitoxin of the escherichia coli RnlA-RnlB toxin-antitoxin module requires RNase HI for inhibition of RnlA Toxin activity. Toxins (Basel). 9, 29 (2017).

54. Goodman, A. L. et al. Extensive personal human gut microbiota culture collections characterized and manipulated in gnotobiotic mice. Proc. Natl. Acad. Sci. U. S. A. 108, 6252–7 (2011).

55. Schnoes, A. M., Brown, S. D., Dodevski, I. & Babbitt, P. C. Annotation Error in Public Databases: Misannotation of Molecular Function in Enzyme Superfamilies. PLoS Comput. Biol. 5, e1000605 (2009).

56. Koonin, E. V. Comparative genomics, minimal gene-sets and the last universal common ancestor. Nat. Rev. Microbiol. 1, 127–136 (2003).

57. Bergmiller, T., Ackermann, M. & Silander, O. K. Patterns of Evolutionary Conservation of Essential Genes Correlate with Their Compensability. PLoS Genet. 8, e1002803 (2012).

58. Patrick, W. M., Quandt, E. M., Swartzlander, D. B. & Matsumura, I. Multicopy Suppression Underpins Metabolic Evolvability. Mol. Biol. Evol. 24, 2716–2722 (2007).

59. Engler, C., Gruetzner, R., Kandzia, R. & Marillonnet, S. Golden Gate Shuffling: A One-Pot DNA Shuffling Method Based on Type IIs Restriction Enzymes. PLoS One 4, e5553 (2009).

60. Salis, H. M. The ribosome binding site calculator. in Methods in Enzymology 498, 19–42 (Academic Press Inc., 2011).

61. Gibson, D. G. et al. Enzymatic assembly of DNA molecules up to several hundred kilobases. Nat. Methods 6, 343–345 (2009).

62. Hartley, J. L., Temple, G. F. & Brasch, M. A. DNA cloning using in vitro site-specific recombination. Genome Res. 10, 1788–95 (2000).

63. Ondov, B. D. et al. Mash: fast genome and metagenome distance estimation using MinHash. Genome Biol. 17, 132 (2016).

64. Steinegger, M. & Söding, J. MMseqs2 enables sensitive protein sequence searching for the analysis of massive data sets. Nat. Biotechnol. 35, 1026–1028 (2017).

65. Steinegger, M. & Söding, J. Clustering huge protein sequence sets in linear time. Nat. Commun. 9, 1–8 (2018).

66. St-Pierre, F. et al. One-step cloning and chromosomal integration of DNA. ACS Synth. Biol. 2, 537–541 (2013).

67. Clermont, O., Christenson, J. K., Denamur, E. & Gordon, D. M. The Clermont Escherichia coli phylo-typing method revisited: improvement of specificity and detection of new phylo-groups. Environ. Microbiol. Rep. 5, 58–65 (2013).

68. Seemann, T. Prokka: rapid prokaryotic genome annotation. Bioinformatics 30, 2068–2069 (2014).

69. Treangen, T. J., Ondov, B. D., Koren, S. & Phillippy, A. M. The Harvest suite for rapid core-genome alignment and visualization of thousands of intraspecific microbial genomes. Genome Biol. 15, 524 (2014).

70. Price, M. N., Dehal, P. S. & Arkin, A. P. FastTree 2 – Approximately Maximum-Likelihood Trees for Large Alignments. PLoS One 5, e9490 (2010).

71. Mitchell, A. L. et al. InterPro in 2019: improving coverage, classification and access to protein sequence annotations. Nucleic Acids Res. 47, D351–D360 (2019).

72. El-Gebali, S. et al. The Pfam protein families database in 2019. Nucleic Acids Res. 47, D427–D432 (2019).

73. Kelley, L. A., Mezulis, S., Yates, C. M., Wass, M. N. & Sternberg, M. J. E. The Phyre2 web portal for protein modeling, prediction and analysis. Nat. Protoc. 10, 845–858 (2015).

74. Deatherage, D. E. & Barrick, J. E. Identification of Mutations in Laboratory-Evolved Microbes from Next-Generation Sequencing Data Using breseq. in Engineering and analyzing multicellular systems 165–188 (Humana Press, New York, NY, 2014). doi: 10.1007/978-1-4939-0554-6_12

75. Langmead, B. & Salzberg, S. L. Fast gapped-read alignment with Bowtie 2. Nat. Methods 9, 357–359 (2012).

76. Li, H. et al. The Sequence Alignment/Map format and SAMtools. Bioinformatics 25, 2078–2079 (2009).

77. Anders, S., Pyl, P. T. & Huber Wolfgang. HTSeq—a Python framework to work with high-throughput sequencing data. Bioinformatics 31, 166–169 (2015).

