## Supplementary Figures for "The impact of genetic diversity on gene essentiality within the *E. coli* species"

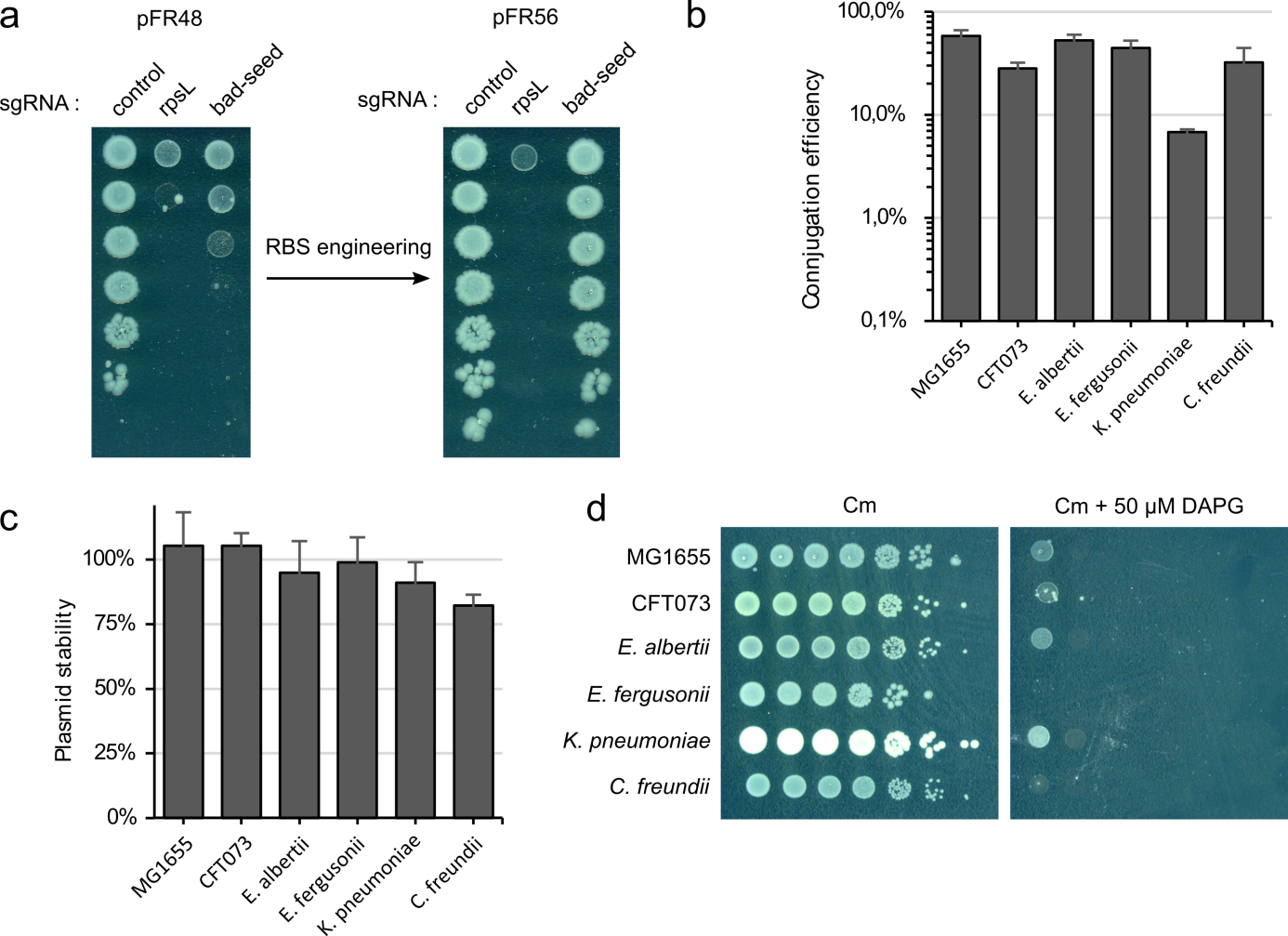

**Supplementary Fig. 1 – The plasmid pFR56 is an optimized and transferrable platform for CRISPRi in *Escherichia* and closely-related genera. a,** The ribosome-binding site of dCas9 was optimized to avoid the previously-described “bad-seed” toxicity effect^1^ while maintaining efficient repression (see Methods). Ten-fold serial dilutions of cells were spotted on LB plates supplemented with chloramphenicol (Cm) and 50 µM DAPG. The nucleotide sequences of sgRNAs are described in Supplementary Table 7. **b**, The conjugation efficiency of pFR56 was assessed by measuring the proportion of chloramphenicol-resistant recipient cells in *E. coli* K-12 MG1655 and CFT073 and in *Escherichia albertii*, *Escherichia fergusonii*, *Klebsiella pneumoniae* and *Citrobacter freundii*. **c,** Plasmid stability was assessed in these strains by measuring the proportion of chloramphenicol-resistant cells after 24 generation of growth in LB without chloramphenicol. **d,** CRISPRi-mediated killing was measured in these strains after conjugation of an sgRNA targeting the essential gene *rpsL*. Ten-fold serial dilutions of cells were spotted on LB plates with chloramphenicol and with or without 50 µM DAPG.

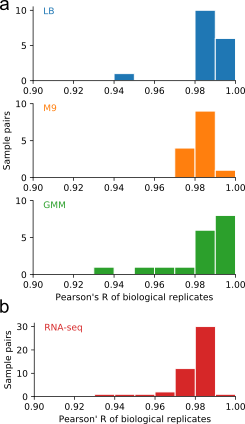

**Supplementary Fig. 2 – CRISPRi screening and RNA-seq experiments are highly reproducible.** (**a-b**) Histograms show the distribution of Pearson’s correlation coefficients between biological replicates. CRISPRi screening experiments were performed in two biological replicates (**a**) with 18 strains in LB (top) and GMM (bottom) and 14 strains in M9-glucose (middle). RNA-seq was performed in three biological replicates in 16 strains (**b**), representing 48 pairwise comparisons.
●

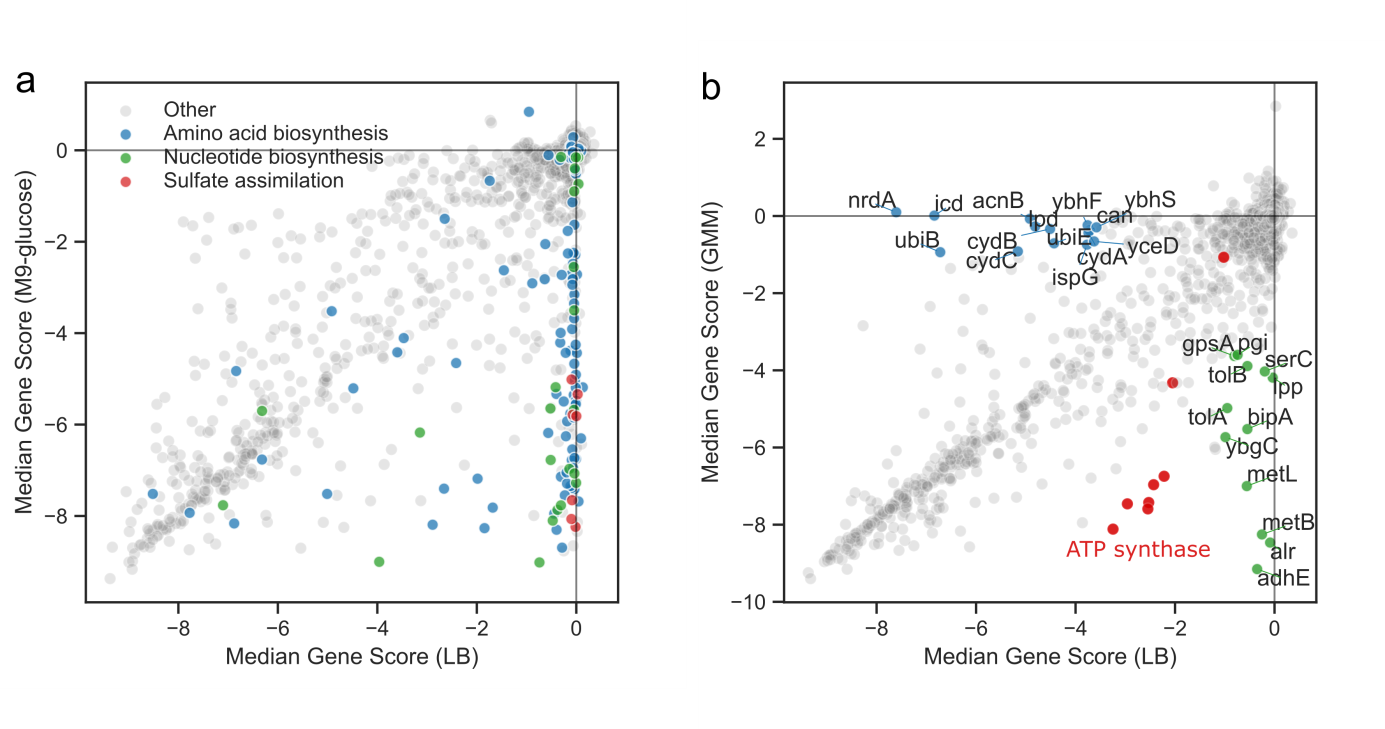

**Supplementary Fig. 3 – Comparison of CRISPRi screening results in different media reveals conditionally-essential genes. (a-b)** The median gene score across all strains was used to assess the most widespread genetic requirements in the *E. coli* species. **a,** Comparison between screening results in LB and M9-glucose medium highlights auxotrophy genes. Genes involved in amino acid and nucleotide biosynthesis and in sulfate assimilation are highlighted. **b,** Comparison of screening results in LB and in GMM reveal medium-specific essential genes. ATP synthase genes are highlighted in red, LB-specific essential genes are highlighted in blue and GMM-specific essential genes are highlighted in green.

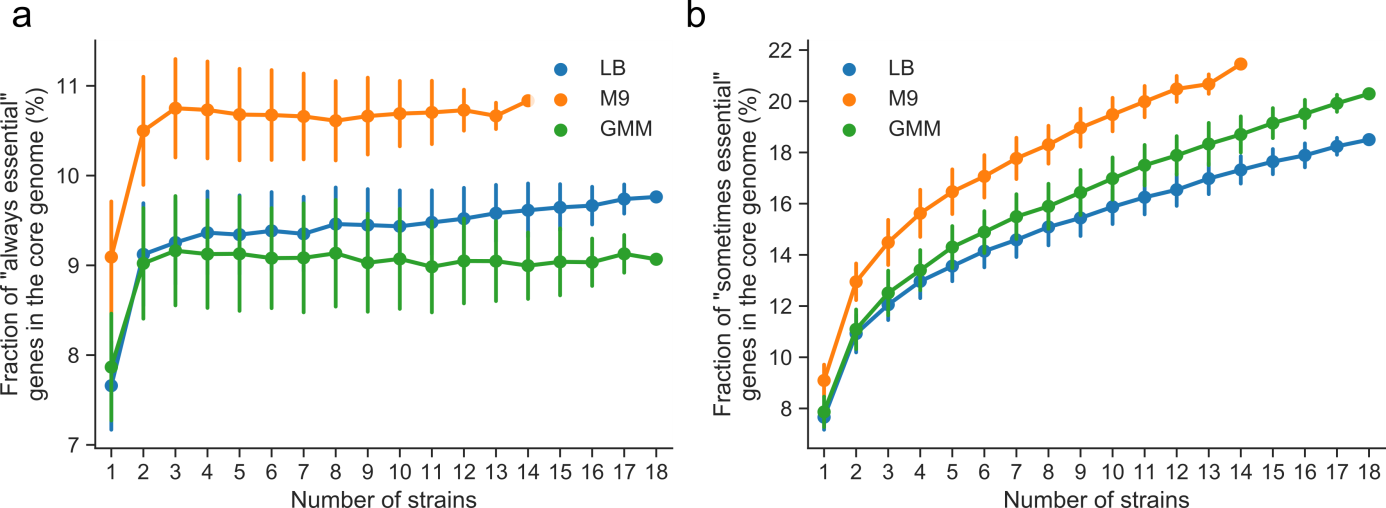

**Supplementary Fig. 4 – Evolution of the proportion of core- and pan-essential genes in the core genome.** The fraction of core genes that are essential in all strains (**a**) or in at least one strain (**b**) was calculated for various sets of strains. Error bars show the standard-deviation of up to 250 permutations.

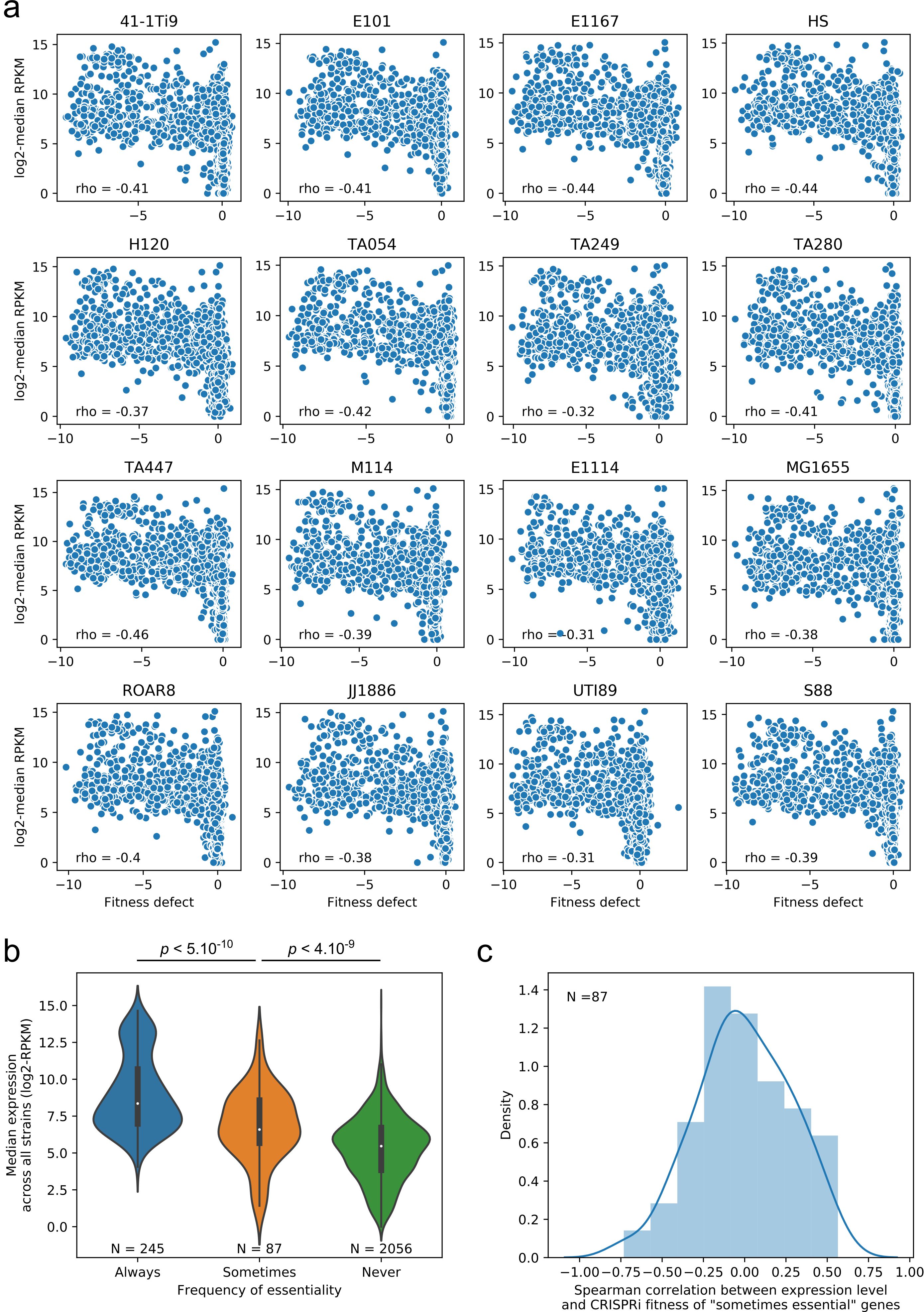

**Supplementary Fig. 5 – The relationship between gene expression and the frequency of essentiality in an *E. coli*** **strain panel. a,** Spearman’scorrelation between gene score in CRISPRi screens and gene expression for 2698 core genes is shown for the 16 strains assayed in both experiments. **b,** We selected 245 genes that are essential in all 16 strains (gene score < -3 in all strains), 87 genes that are variably essential (gene score < -3 in ≥1 strain and gene score > -1 in ≥1 strain), and 2056 genes that are never essential in the tested strains (gene score > -1 in all strains). Violin-plots show the distribution of the median gene expression level across all strains. Inside each distribution, the white dot shows the median and the extremities of the black bar show the 1^st^ and 3^rd^ quartiles of the distribution. p-values were calculated using a two-sided Mann-Whitney U test. **c,** For each “variably essential” gene, we calculated Spearman’s correlation coefficient between gene expression level and gene score across the 16 strains. This plot shows the distribution of these correlation coefficients.

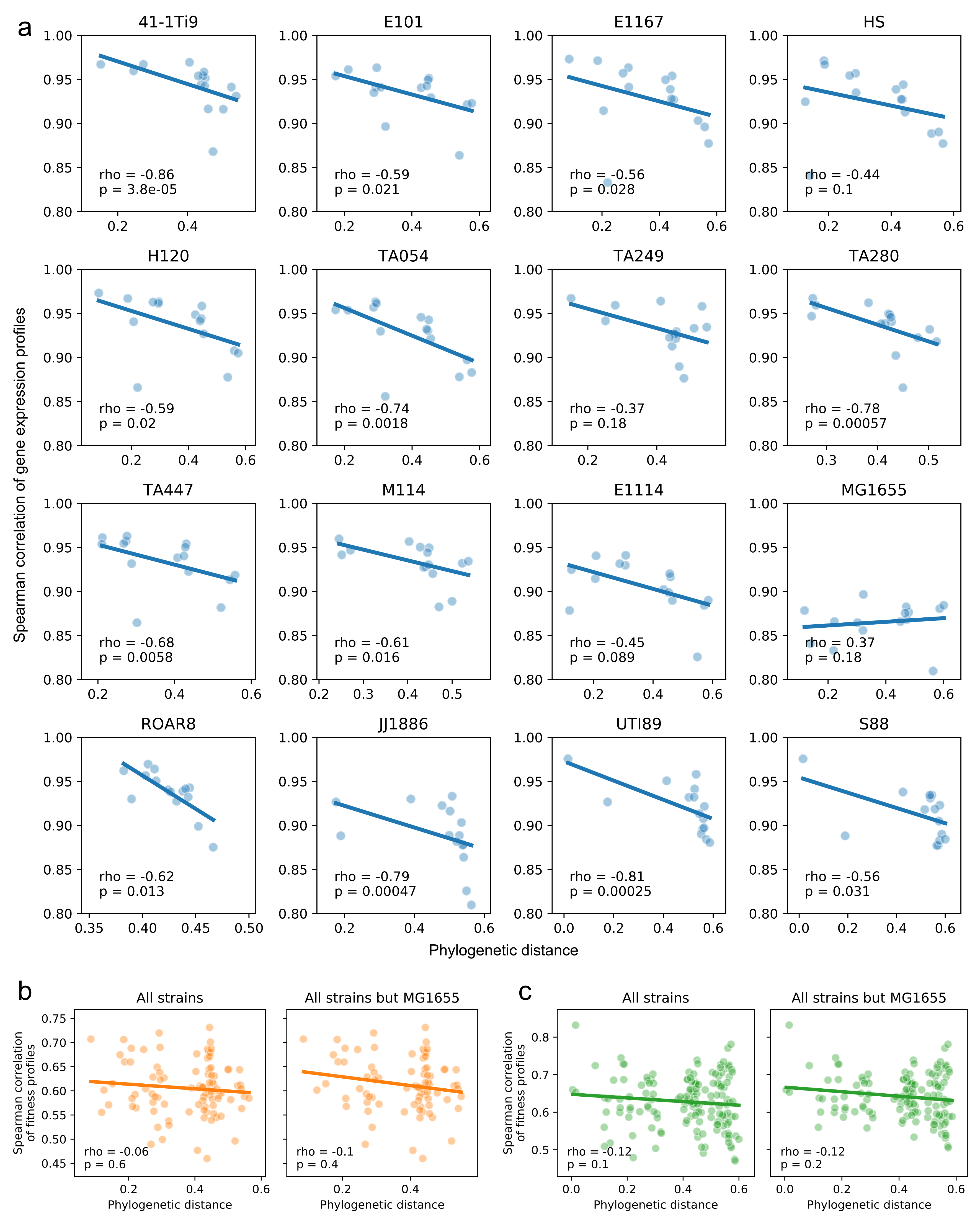

**Supplementary Fig. 6 – The influence of phylogeny on gene expression and gene essentiality.** (**a-c**) For each pair of strains, we used Spearman’s correlation coefficient as a measure of the similarity in gene expression profile or gene essentiality profile. **a,** Each plot represents the relationship between the phylogenetic distance and the similarity in gene expression profile between a given strain and the 15 remaining strains assayed during RNA-seq experiments. **(b-c)** There is no significant correlation between the phylogenetic distance of pairs of strains and their similarity in gene essentiality profiles in M9 (**b**) and in GMM (**c**). Including (left) or excluding (right) MG1655 in this analysis had no impact.

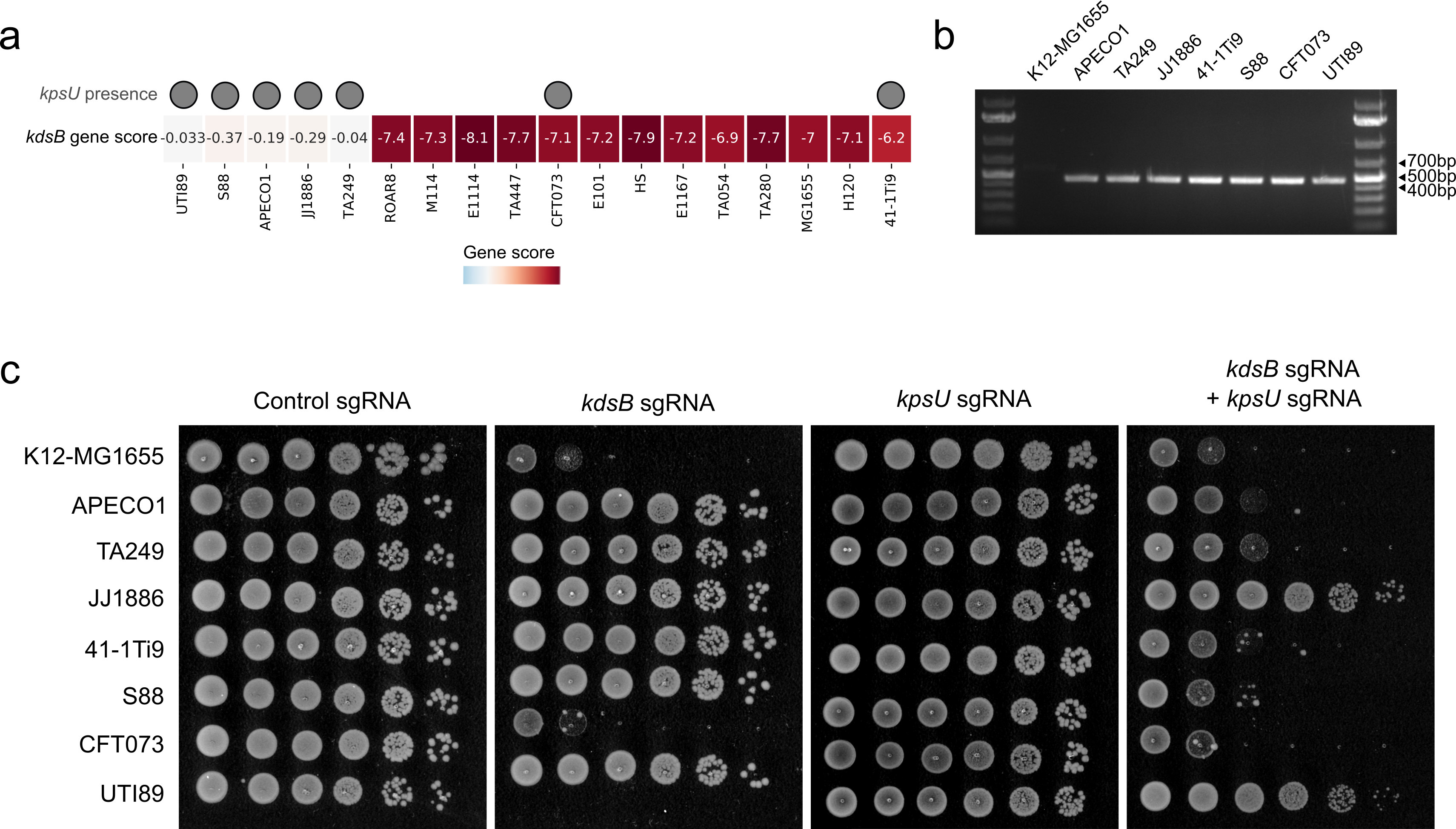

**Supplementary Fig. 7 – Functional redundancy between *kdsB* and *kpsU*. a**, A heatmap shows the fitness defect associated with the repression of *kdsB*. The gene score values are shown inside each box. Grey circles represent the presence of *kpsU* in the genome. **b**, A PCR using *kpsU-*specific primers (FR257 and FR258, **Supplementary Table 6**) followed by gel electrophoresis shows the presence of *kpsU* in selected strains. A 469-bp product is expected when *kpsU* is present. **c**, Spot assays show the phenotype associated with the repression of *kdsB*, *kpsU* or both *kdsB* and *kpsU* simultaneously in strains carrying *kpsU*. K-12-MG1655 does not have a copy of *kpsU* and was used as a control. Ten-fold serial dilutions were plated on LB supplemented with chloramphenicol and 50 µM DAPG. The nucleotide sequences of sgRNAs are shown in **Supplementary Table 7**.

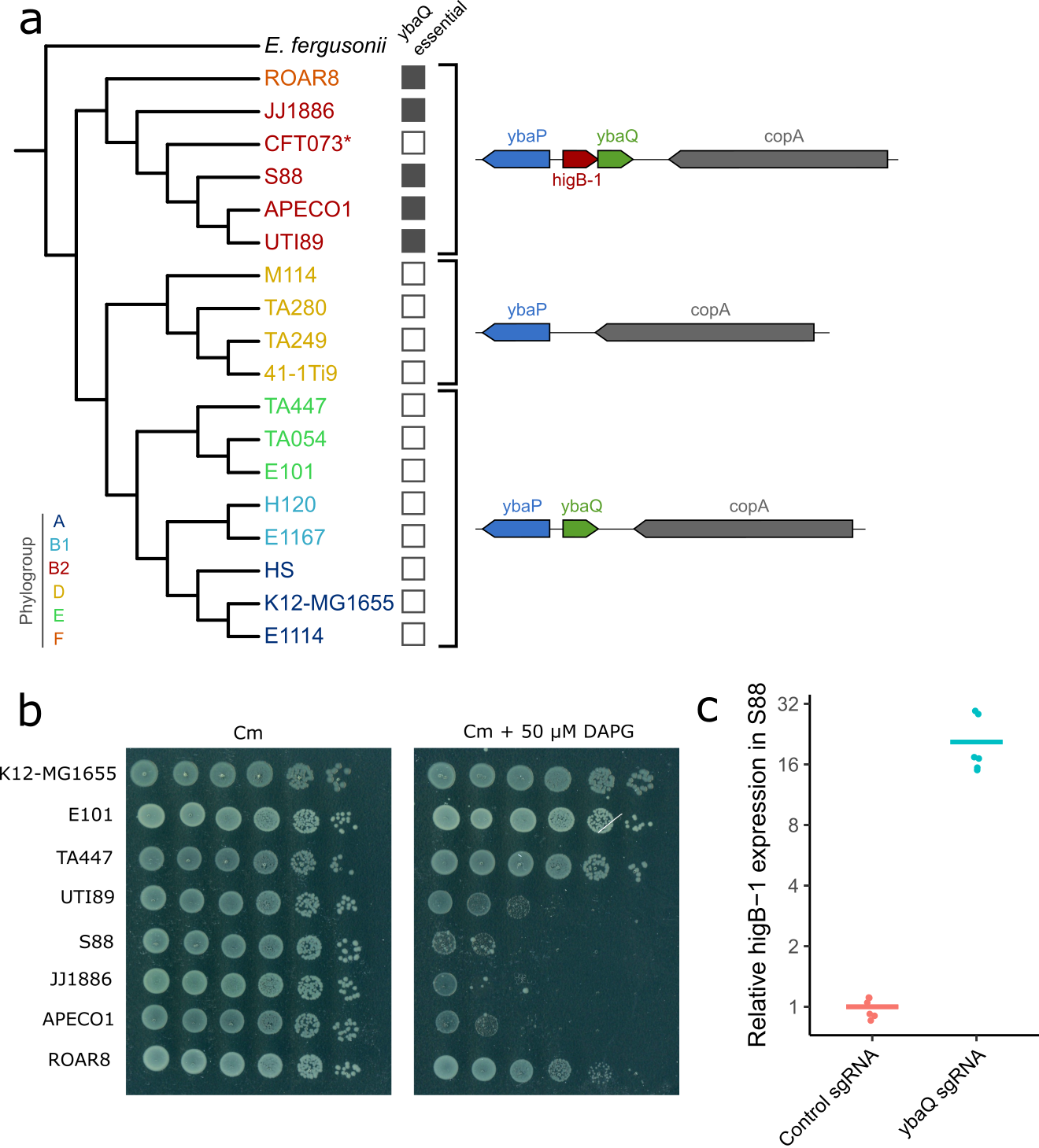

**Supplementary Fig. 8 – *ybaQ* is a transcriptional repressor of the HigB-1 toxin. a,** A phylogenetic tree was built with the 18 strains assayed in CRISPRi screens. Grey squares indicate if *ybaQ* is essential in the corresponding strains. The genomic region of the *ybaQ* locus is shown on the right and varies according to three clades. (*) *higB* is interrupted by a stop codon in strain CFT073. **b,** CRISPRi screens results were validated by introducing *ybaQ* sgRNA in three strains where *ybaQ* repression has no effect (K12-MG1655, E101 and TA447) and in five strains where *ybaQ* repression is toxic (UTI89, S88, JJ1886, APECO1 and ROAR8). Ten-fold serial dilutions of cells were spotted on LB plates containing chloramphenicol with or without 50 µM DAPG to induce dCas9 expression. The sequence of the *ybaQ* sgRNA is provided in **Supplementary Table 7**. **c,** Silencing *ybaQ* in strain S88 induces the expression of *higB-1*. The horizontal bar shows the mean of 3 biological replicates and 2 technical replicates from RT-qPCR experiments. Primers used for qPCR are shown in **Supplementary Table 8**.

**Supplementary references**

1.	Cui, L. *et al.* A CRISPRi screen in E. coli reveals sequence-specific toxicity of dCas9. *Nat. Commun.* **9**, 1912 (2018).
